# Decoding single-cell replication states from imaging data reveals locus-specific subnuclear position dependencies of replication initiation

**DOI:** 10.64898/2026.08.12.744402

**Authors:** Francesco Musella, David M. Gilbert, Frank Alber

**Author notes:** Correspondence: David M. Gilbert, Frank Alber.

## Abstract

Replication timing (RT) is a fundamental feature of genome regulation, tightly linked to chromatin state and three-dimensional (3D) nuclear organization. Yet how the reproducible population-level timing program emerges from stochastic replication decisions in individual cells remains unclear, because it requires measuring RT and 3D genome organization jointly in the same cell.

Here, we introduce RepTile, the first systematic framework that infers allele-resolved, single-cell replication states directly from multiplexed FISH and chromosome tracing experiments. By extracting replication information from spot-count distributions while correcting for genomic and spatial detection biases, RepTile infers cell-cycle progression, locus-specific replication timing, cell-to-cell heterogeneity in replication timing, and replication states of individual allele-resolved locus copies within their native three-dimensional nuclear context.

Using RepTile on mouse embryonic stem cells, we find that the association between replication timing and subnuclear position observed in population-level genomic experiments conceals fundamentally different locus-specific behaviors. We found two classes of chromatin regions: “position-sensitive” and “position-insensitive”. Position-sensitive regions replicate at different times depending on their subnuclear location, typically firing earliest when positioned in their canonical locale. Strikingly, this holds even when the canonical locale is repressive, such as the nuclear envelope. On the other hand, position-insensitive loci replicate at their characteristic times regardless of subnuclear position.

These classes map onto distinct regulatory regimes, which are particularly pronounced at replication initiation zones (IZ): position-sensitive IZs are enriched in domains that replicate constitutively early or late across cell types, whereas position-insensitive IZs are enriched in domains that switch RT developmentally and contain early replication control elements (ERCEs), cis-regulatory elements known to govern IZ firing times.

Together, these results challenge the view that subnuclear location broadly defines replication timing, indicating that positional dependence is specific to constitutive domains, while developmental domains carry intrinsic cis-acting programs that decouple their replication from subnuclear location.

## Introduction

Genome organization and replication timing are coordinately re-established in early G1 at the timing decision point (TDP), raising the long-standing question of how chromatin folding, subnuclear positioning and DNA replication are related within individual cells ^1–7^. Resolving these relationships requires measurements of both the spatial organization and replication state of genomic loci in the same cell. Sequential chromosome tracing and multiplexed spatial-omics fluorescence in-situ hybridization (FISH) methods ^8–17^ now enable the locations of hundreds to thousands of genomic loci to be measured in individual cells, in some cases at whole-genome scale. Approaches such as DNA MERFISH ^13^, DNAseqFISH+ ^16,18,19^, and ORCA ^11^ use iterative rounds of oligonucleotide hybridization, fluorescence imaging and signal removal, with locus identities encoded through sequential or combinatorial labeling schemes. Many platforms can additionally map RNA transcripts and nuclear structures in the same cells, providing a multimodal view of nuclear genome organization within its transcriptional and subnuclear context ^13,16,20^. These capabilities make it possible to systematically investigate how chromatin folding and nuclear positioning relate to other nuclear processes.

However, these methods currently lack a systematic framework for inferring locus-resolved replication states in proliferating cells. In conventional DNA FISH, replication has often been inferred from closely spaced signal doublets or, when replicated copies remain spatially unresolved, from increased fluorescence intensity of a single signal ^21–24^. However, such heuristic assignments are difficult to apply to modern chromosome-tracing and multiplexed FISH experiments, in which loci are commonly targeted by large numbers of tiled primary probes to increase detection sensitivity and decoded across sequential imaging rounds ^8,25^. An unreplicated locus may therefore appear as multiple spatially separated spots, whereas incomplete hybridization, decoding errors or missed detections can cause a replicated locus to yield only one or no detectable signal. Consequently, observed spot counts cannot be translated directly into DNA copy number. Thus, despite the long-standing use of conventional FISH to assess replication at individual loci, a systematic framework for inferring single-cell, locus-and allele-resolved replication states from multiplexed spatial-genomics data has so far been lacking.

Resolving replication state is also important for the quantitative interpretation of multiplexed imaging data, because latent copy-number variation can bias measurements of locus-locus distances, chromatin compactness or chromosome-territory volumes. Beyond correcting these biases, resolving replication states within their native nuclear context would enable DNA replication timing to be studied jointly with chromatin folding, subnuclear positioning and gene activity in the same cell. Such measurements have not previously been possible and would enable a direct means of determining whether nuclear position is functionally coupled to replication timing and whether this relationship applies uniformly across the genome.

For these reasons, we developed RepTile, a computational framework that infers cell-cycle progression and locus-resolved single-cell replication states directly from multiplexed FISH and chromosome tracing data. RepTile treats replication state as a latent variable and infers replication information from noisy per-locus spot-count distributions while explicitly accounting for detection efficiency, signal overcounting, and spatial imaging biases. By combining statistical inference with machine-learning, RepTile reconstructs genome-wide replication timing and its heterogeneity, S-phase progression and the replication states of individual, spatially resolved locus copies within their native three-dimensional nuclear context in single cells. This enables replication dynamics to be analyzed together with chromatin folding, subnuclear position and nuclear environment across single cells.

Using RepTile, we show that the previously observed population-level association between replication timing and subnuclear position ^1,5,7,26,27^ conceals fundamentally different behaviors among genomic loci. Replication at some loci is highly position sensitive: their probability of replication changes markedly depending on the nuclear locale they occupy. The same nuclear environment can advance replication for one class of loci while delaying it for another. A common principle, however, is that position-sensitive loci generally replicate earliest in their preferred nuclear environment, whether interior active, nucleolar or peripheral repressive. In contrast, other loci remain largely position insensitive, replicating at similar times across distinct nuclear environments despite retaining strong spatial preferences in their nuclear locations.

Moreover, at initiation zones, the distinction between position-sensitive and position-insensitive loci was closely linked to developmental replication-timing plasticity: strongly position-sensitive initiation zones preferentially occur within domains that maintain constitutively early and late replication across cell types, whereas position-insensitive initiation zones preferentially occur within developmentally regulated domains that switch replication timing during differentiation. Together, these findings reveal distinct position-dependent and position-insensitive, likely locus-intrinsic modes of replication control and challenge the view that subnuclear position exerts a uniform influence on replication timing across the genome. Our results therefore refine the long-standing model emerging from the TDP: although nuclear organization and replication timing are coordinately established, the extent to which replication timing depends on nuclear position differs substantially among genomic loci and is linked to developmental replication-timing plasticity.

## Results

### RepTile: a multi-modal framework to infer replication from Multiplex FISH data

RepTile is a multi-modal statistical and machine learning framework that infers cell-cycle stage and locus-resolved single-cell replication states from genome-wide, multiplex FISH data (**Figure 1a**). RepTile first parses the imaging data to extract the spot count per locus allele, i.e., the number of spot detections assigned to each allele-resolved locus in each cell, together with auxiliary features, including per-spot fluorescence intensity and 3D positions of alleles relative to imaged nuclear landmarks, including nuclear speckles, nucleoli and the nuclear envelope (**Methods**). Although image processing, spot calling and barcode decoding differ across M-FISH platforms ^28^, RepTile operates downstream of these steps and requires only a decoded table of locus-associated spot detections, making it broadly applicable across imaging platforms. It leverages two complementary modules to infer replication dynamics: (i) a statistical module, and (ii) a machine-learning module.

**Figure 1.**
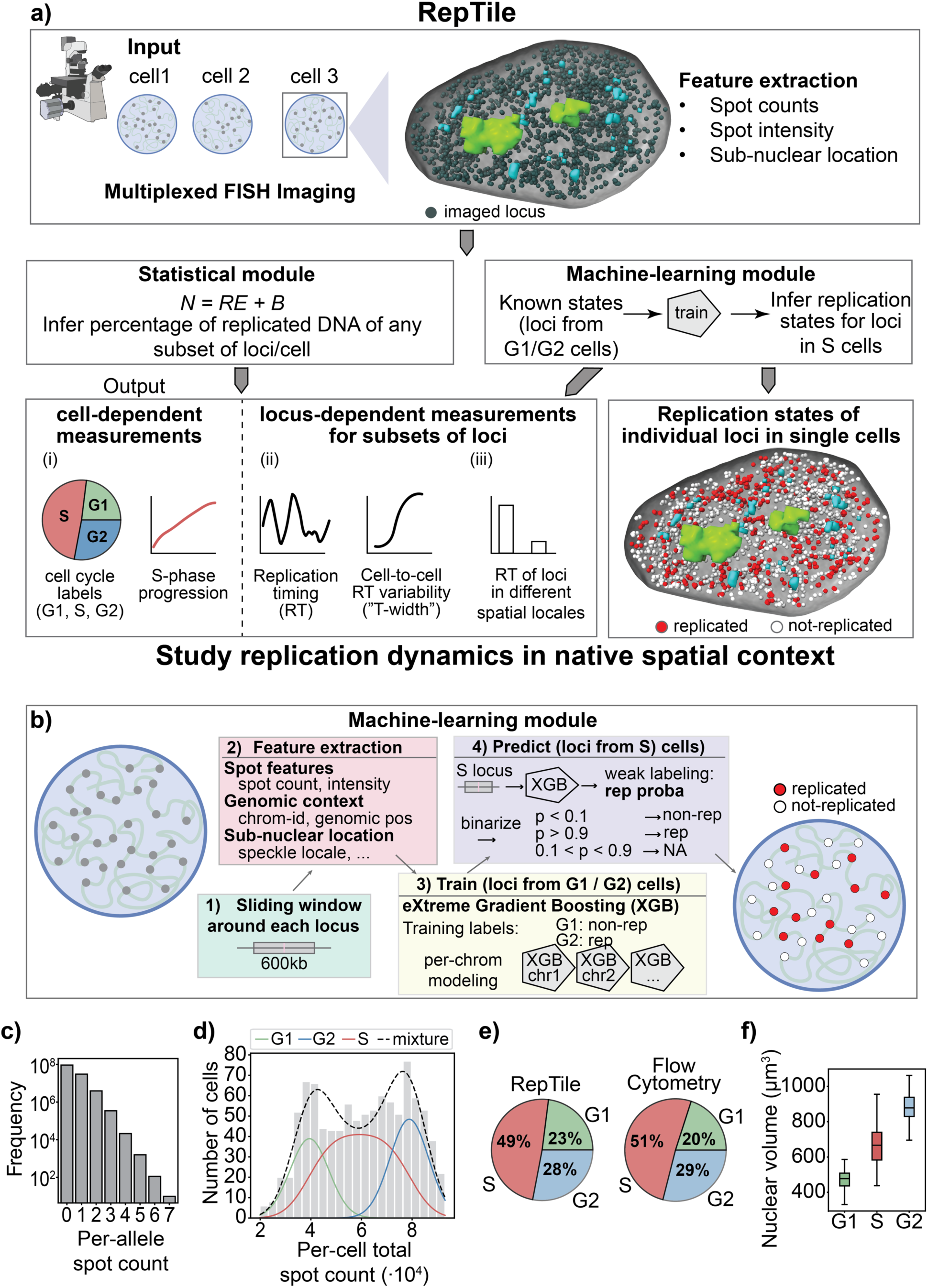
Schematic overview of RepTile for identifying cell-cycle progression and single-cell replication states from multiplexed FISH and chromosome tracing data. **a)** Schematic flowchart of RepTile. Top panel: Imaging data is parsed into spatial features used for cell-cycle and DNA replication inference. The main feature is the ‘spot count’, defined as the number of spatially resolved imaged signals per locus in each cell. An example DNAseqFISH+ cell is shown ^16^. Middle panels: RepTile contains two complementary modules. Left middle panel: the statistical module infers the percentage of replicated DNA from any subset of loci and cells, allowing us to measure cell-specific S-phase progression and locus-dependent replication properties. Right middle panel: the machine-learning module uses loci in G1-and G2-phase cells to train eXtreme Gradient Boosting (XGB) models ^30^ to predict the replication state of each locus’s allele copy in each single S-phase cell. The same cell shown at the top is displayed with imputed locus-specific replicated states. **b)** Overview of the machine-learning module: features are extracted from a sliding genomic window centered at each locus. Extreme gradient-boosting (XGB) models are trained for each chromosome using loci from G1-and G2-phase cells. The trained models are applied to S-phase cells and replication states of loci are predicted by binarizing the weak labeling from XGBs (**Methods**). **c)** Spot count distribution per locus allele for the E14 mESC 2025 dataset ^16^. **d)** Distribution of per-cell total spot counts. Cells were assigned to G1, S or G2 using a mixture-model fit (**Methods**). Green, red and blue curves indicate the predicted G1, S, G2 distributions, respectively, and the dashed black curve indicates the combined fitted distribution. **e)** Percentage of cells assigned to G1, S and G2 by RepTile (left) and measured in an independent flow-cytometry experiment (right) ^33^. **f)** Box-plot distributions of single-cell nuclear volumes for G1, S and G2 cells.

The statistical module estimates the fraction of replicated DNA for any chosen subset of loci and cells in the dataset. We model the observed spot count *N*_*ic*_ for locus *i* in cell *c* as *N*_*ic*_ = *R*_*ic*_*E*_*ic*_ + *B*_*ic*_, where *R*_*ic*_ ∈ {1,2} represents the replication state (non-replicated vs. replicated), *E*_*ic*_ is a detection factor modeled by the detection efficiency *ε*_*ic*_, and *B*_*ic*_captures background and false positives with an overcounting rate *β*_*ic*_ (**Methods**). Since the replication state is known in G1 and G2 stages, we first calibrate the detection parameters (*ε*_*ic*_, *β*_*ic*_) using G1 and G2 cells, and then hold these parameters fixed to infer the replicated fraction in S-phase for any subset of loci and cells (**Methods**). These quantities are estimated via the Generalized Method of Moments by matching empirical moments of the observed spot-count distribution to the corresponding moments predicted by our parametric model, yielding a system of equations for the unknowns to be solved. This yields (**Figure 1a**): (i) cell-dependent estimates of cell-cycle stage and S-phase progression; (ii) locus-dependent estimates of replication timing and its cell-to-cell variability; and (iii) spatially dependent estimates linking replication to sub-nuclear locations and compartments. G1/S/G2 cell labels can either be provided by orthogonal methods such as DAPI ^13,29^ or, as shown in the next section, inferred by RepTile itself. RepTile’s machine learning module predicts the single-cell replication state of each allele-resolved locus (replicated vs. non-replicated) (**Figure 1ab**, **Methods**). For each locus, we extract features from a sliding window over its local neighborhood, including spot count, fluorescence intensity, and genomic and spatial context. We then train chromosome-specific eXtreme Gradient Boosting (XGB) classifiers ^30^ using loci from G1 and G2 cells, labeling loci in G1 cells as non-replicated and those in G2 as replicated. This training also allows the model to learn platform-specific detection biases. Finally, we apply the trained classifiers to S-phase cells to predict locus replication states.

We applied RepTile to a DNAseqFISH+ dataset from E14 mouse embryonic stem cells (mESC) ^16^ with consecutively tiled 25kb regions imaged genome-wide in 994 cells (**Methods**).

### Assigning single cells to G1, S, and G2

We first assigned cells to G1, S or G2 phase to determine their cell-cycle stage. Although the spot counts varied substantially among loci across the genome (**Figure 1c**), the total number of detected spots per cell showed two distinct peaks connected by a broad distribution of intermediate values (**Figure 1d**). This profile resembled the DNA-content distributions obtained by fluorescence-activated cell sorting (FACS) ^31,32^, in which the two peaks correspond to G1 and G2 cells with 2C and 4C DNA content, respectively, whereas cells with intermediate content are undergoing DNA replication in S-phase.

To assign cell-cycle states from this distribution, we fitted a three-component mixture model. G1 and G2 cells were represented by Gaussian distributions centered on the expected 2C and 4C DNA-content levels, respectively, whereas S-phase cells were modeled as the convolution of a uniform distribution, representing the progressive increase in DNA content during replication, with a Gaussian distribution accounting for the cell-to-cell variation in spot count detection (**Figure 1d**, **Methods**). Our model provided a good fit to the data (Kolmogorov-Smirnov goodness-of-fit test, p = 0.63), and classified 226 cells as G1 (22.7%), 490 as S (49.3%) and 278 as G2 (28.0%).

The inferred cell-cycle fractions closely agreed with those measured independently by flow cytometry in the same cell line ^33^, differing by an average of only 1.8% across the three phases (**Figure 1e**). Consistent with the assigned cell-cycle states, nuclear volume increased progressively from G1 through S to G2 (**Figure 1f**), as expected for proliferating cells.

Finally, we assessed the robustness of the cell-cycle assignments using two independent classification strategies based on orthogonal information: a two-component Gaussian mixture model fitted to the spot counts of the earliest-and latest-replicating loci, and a nuclear-volume-based ranking calibrated to the cell-cycle fractions measured by FACS (**Methods, Extended Figure 1abc**). Both approaches showed strong agreement with the primary classification, with a mean agreement of 85.5% (**Extended Figure 1d**), indicating that the inferred assignments were not dependent on a single modeling strategy.

### Chromatin organization is established by S phase and largely retained thereafter

Proximity-frequency maps derived from imaging data showed strong agreement with Hi-C ^34^ for both intra-chromosomal (250kb: Pearson r = 0.91, p-value < 1e-300, adjusted Pearson r = 0.63) and inter-chromosomal proximities (1Mb: Pearson r=0.73, p-value < 4.1e-37, **Extended Figure 2ab, Methods**). Contact-proximity maps stratified by cell cycle phase (**Figure 2ab**) showed fewer short-range (< 10Mb) and more long-range (>10 Mb) proximities in G1 than in S and G2 cells, consistent with reduced local compaction and increased large-scale chromatin mixing (**Figure 2a**). These changes coincided with a strengthening of compartments from G1 to S, as shown by higher compartment scores in S than in G1, confirming recent Hi-C studies ^35^ (**Figure 2c**, **Extended Figure 2c**, **Methods**). This strengthening was driven primarily by a pronounced gain of A–A contacts in S phase relative to G1, followed by a partial reduction in G2 (**Figure 2de**).

**Figure 2.**
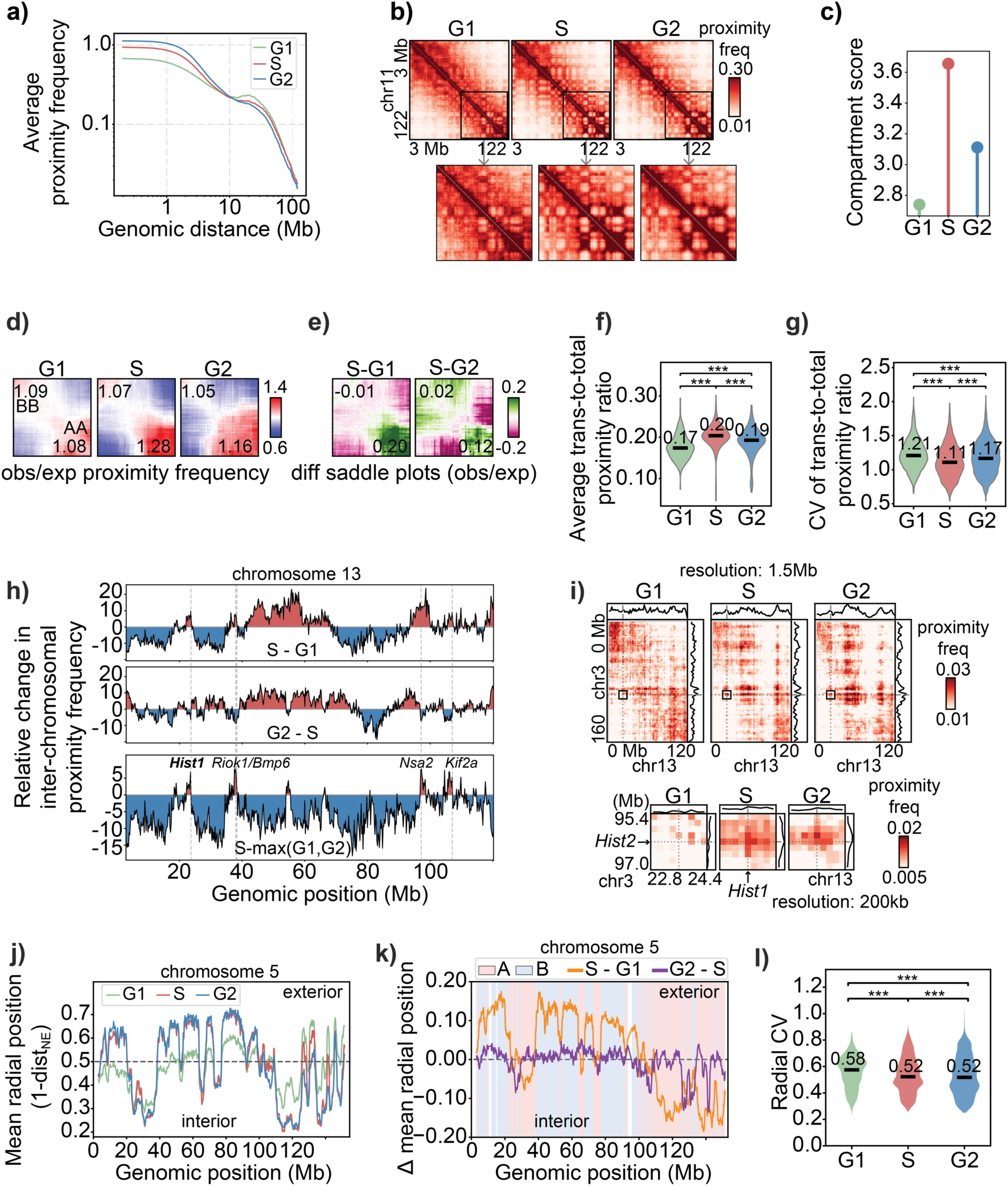
Multi-scale re-structuring of chromatin throughout the cell-cycle revealed by RepTile’s single-cell G1/S/G2 assignment. **a)** Average proximity frequency as a function of genomic distance in G1, S, G2-phase cells (**Methods**). **b)** Intrachromosomal proximity frequency maps averaged across G1 (left), S (middle) and G2 (right) cells for chromosome 11, with a zoomed view of a 50Mb region. **c)** Average compartment score calculated in G1, S and G2 cells (**Methods**). **d)** Saddle plots displaying the average observed-to-expected proximity frequencies between loci stratified and ordered by their PC1 compartment score, revealing preferential A-A, B-B and A-B contacts (**Methods**). The top-left corner corresponds to B-B interactions, the bottom right to A-A, and the anti-diagonal shows inter-compartment interactions. The numbers report the top A-A and top B-B interactions. **e)** Difference saddle plots comparing S with G1 (S-G1) and S with G2 (S-G2). **f-g)** Violin plots showing the distributions of the average trans-to-total ratio (**f**) and its coefficient of variation (CV; **g**) across G1-, S-, G2-phase cells (**Methods**). Three stars indicate significance by two-sided Mann-Whitney U (f) or permutation test (g) (p < 0.001, **Methods**). **h)** Relative percentage change in the row sum of inter-chromosomal proximities for each locus on chromosome 13 across all interactions with loci on other chromosomes (*τ*_*i*_, **Methods**). Changes are shown for S relative to G1 (100 (*τ*_*i*,*S*_ − *τ*_*i*,*G*1_)⁄*τ*_*i*,*G*1_) (top panel), G2 relative to S, 100 (*τ*_*i*,*G*2_ − *τ*_*i*,*S*_)⁄*τ*_*i*,*S*_ (middle panel), and S relative to max(G1,G2), 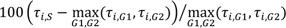 (bottom panel). Vertical dashed lines mark selected genes and genomic regions showing S-phase specific increases in chromosomal proximities, including the *Hist1* gene cluster. **i)** Inter-chromosomal proximity frequency maps between chromosomes 3 and 13 in G1, S, G2 at 1.5Mb resolution (top) and a 200kb resolution zoom of a 1.8Mb region spanning the *Hist1*-*Hist2* gene cluster pair (bottom, **Methods**). **j-k)** Average radial position for loci along chromosome 5 (**Methods**) at 200kb resolution for G1-, S-, and G2-phase cells (**j**) and the corresponding S-G1 and G2-S differences (**k**). In **k**, red and blue shaded regions correspond to A and B compartments, defined from Hi-C data ^37^. **l)** Violin plots showing the distributions of the radial position coefficient of variation (CV) across G1-, S-, G2-phase cells. The three stars indicate significance by paired Wilcoxon test (p-value < 0.001).

The increased compartmental segregation in S-phase cells was accompanied by increased inter-chromosomal proximities across the genome. The mean trans-to-total proximity frequency ratio was significantly higher in S phase than in G1 (Mann-Whitney p = 1.0e-36) and, to a lesser extent, G2 (Mann-Whitney p = 4.3e-11, **Figure 2f**, **Methods**). Moreover, G1 cells showed a greater cell-to-cell variability in chromosome folding, reflected by a higher coefficient of variation in the trans-to-total proximity ratio than in S or G2 cells (two-sided permutation test p-values ≤ 4.0e-4, **Figure 2g**, **Methods**). We further identified 1,558 genomic regions at 200kb resolution that showed an S-phase specific increase in the trans-to-total contact proximity frequency ratio relative to both G1 and G2 (Δ = S – max(G1, G2), two-sided permutation test Benjamini-Hochberg FDR < 0.05, **Methods**). Chromosome 13, for example, contains several regions with S-phase specific increases in inter-chromosomal proximity frequency (**Figure 2h**, **Methods**). These included the *Hist1* gene cluster (chr13:21.6-24.0Mb), which contains multiple replication-dependent histone genes, and other genes involved in cell cycle regulation and growth (e.g., Riok1, Bmp6, Nsa2, Kif2a). Notably, compared to G1, S-phase cells showed significantly increased proximity between the Hist1 gene cluster on chromosome 13 and the Hist2 gene cluster on chromosome 3, which contains additional replication-dependent histone genes (z-test z = 3.5, p = 4.5e-4, **Figure 2i**, **Extended Figure 2d**, **Methods**). Similar S-phase-specific inter-chromosomal clustering of replication-dependent histone genes has been reported during the early cleavage stage of sea urchin embryos ^36^, but, to our knowledge, has not previously been reported in vertebrate cells. Finally, we examined radial genome organization across the cell cycle, by quantile-normalizing each locus’s distance to the nuclear envelope to account for nuclear expansion (**Methods**). Genome-wide mean envelope distances correlated strongly with Lamin B1 TSA-seq signal ^27^ (Pearson r = 0.92, p < 1e-300), validating our NE distance measure. Radial organization changed primarily between G1 and S (**Figure 2jk**): 70.6% of loci differed significantly in mean NE distance, compared with 23.9% between S and G2 (per-locus Mann–Whitney U, FDR < 0.05, **Methods**). G1 cells showed a narrower range of mean NE distances (**Figure 2j**), combined with greater cell-to-cell positional heterogeneity (paired Wilcoxon G1 > S p = 2.9e-243, G1 > G2 p = 8.8e-217, **Figure 2l**). Positional heterogeneity decreased in S, accompanied by stronger radial segregation of A and B compartment regions. Loci shifting inward were predominantly (90%) A compartment, whereas those positioned closer to the envelope in S belonged overwhelmingly to the B compartment (88%). Thus, positional changes positively correlated with Hi-C compartment PC1 (*r* = 0.72, *p* < 1e-300) and SON TSA-seq (*r* = 0.77, *p* < 1e-300) and anticorrelated with Lamin B1 TSA-seq (*r* = −0.77; *p* < 1e-300). In contrast, S-to-G2 repositioning showed little compartment association (PC1, *r* = 0.17, p-value = 2e-75), indicating that radial segregation of active and repressive chromatin is largely established by S.

Together, these analyses reveal a coordinated reorganization of the genome as cells enter S phase. These findings support a model in which G1 nuclei occupy a relatively heterogeneous and weakly segregated organizational state, whereas entry into S phase establishes a more reproducible and functionally structured genome architecture. This may partly reflect the unusually short G1 phase of mESC cells, which could limit the time available for genome organization to re-establish a stable interphase configuration after mitosis. The comparatively modest changes between S and G2 further indicate that much of this organization is established by S phase and largely retained thereafter.

### Measuring replication timing by RepTile

Next, we used RepTile to infer the replication timing (RT) of loci, defined as the average point in S phase at which a locus replicates ^32^. To achieve this, we first estimated locus-specific detection efficiencies in G1 and G2 cells, where the replication state is known (**Methods**). These estimates were highly correlated between G1 and G2 cells (Pearson r = 0.83, p-value < 1e-300), allowing detection efficiencies in S-phase to be approximated by their mean (**Extended Figure 3ab**).

Detection efficiencies varied systematically across genomic and spatial contexts. They were higher for loci in the inactive B-compartment than in the active A compartment ^37^ (Spearman r_S_ = -0.51, p-value < 1e-300, **Extended Figure 3c**), decreased with axial distance from the microscope lens, and were higher near the nuclear envelope than near speckles or nucleoli (**Extended Figure 3d**). These systematic biases underscore the need to model detection efficiencies when inferring replication dynamics.

Using these locus-specific detection efficiencies, RepTile inferred the replicated fraction of each locus across S-phase cells while accounting for overcounting (**Methods**, **Supplementary Figure 1**). Replication timing was defined as the percentage of S-phase allele instances in which a locus was inferred to be replicated (**Methods**). The resulting genome-wide RT profile closely matched independent ensemble Repli-seq ^38^ (**Figure 3a**) and single-cell Repli-seq ^39^ experiments (**Extended Figure 3e**) with high correlation (Pearson *r* = 0.92 and *r* = 0.88, respectively; both p-value < 1e-300). By contrast, estimating RT directly from uncorrected spot counts substantially reduced agreement with ensemble Repli-seq (Pearson *r* = 0.28, **Methods**), demonstrating that correction for locus-specific detection efficiency is essential for accurate RT inference.

**Figure 3.**
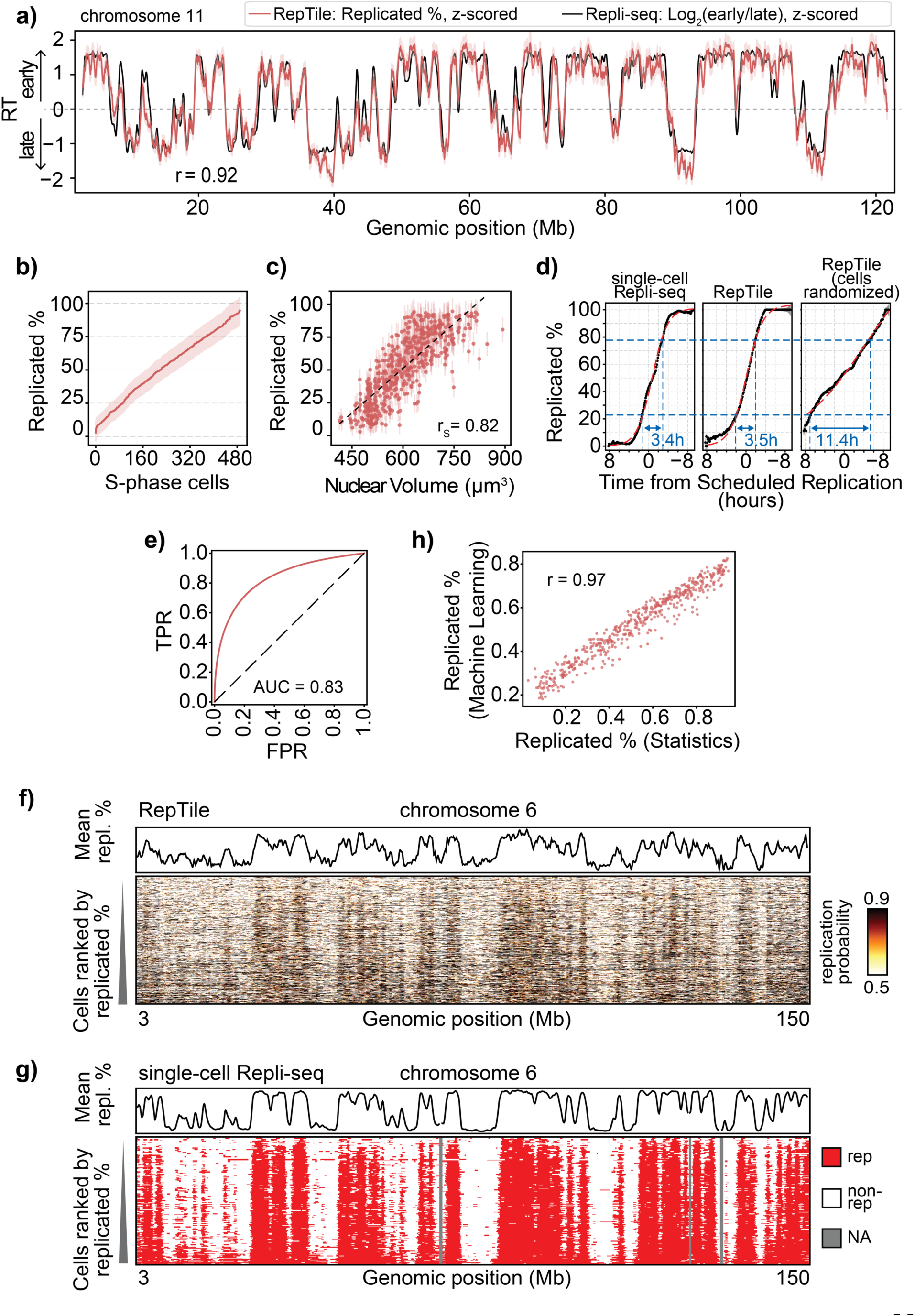
RepTile uncovers single-cell, allele-resolved replication dynamics from DNAseqFISH+ imaging. **a)** The Replication Timing (RT) from Repli-seq (black) ^38^ is shown together with the inferred RT from RepTile’s statistical module (red, **Methods**) for chromosome 11. Both signals were z-scored for visual comparison. The Pearson correlation between them is shown. **b)** Percentage of replicated DNA inferred for individual S-phase cells by the RepTile statistical module (**Methods**). Cells are ordered by their inferred replicated DNA percentage. Error uncertainties are shown as a shaded area. **c)** Scatter plot of inferred percentage of replicated DNA against the corresponding nuclear volume in individual S-phase cells. Vertical lines indicate error uncertainties in the inferred replicated %, and the dashed black line shows the linear regression fit. The Spearman correlation is shown. **d)** Percentage of replicated loci, averaged across all S-phase cells, against Time from Scheduled Replication, binned every 0.1 hours, calculated from single-cell Repli-seq ^39^ (left), inferred from RepTile (middle) and obtained from RepTile after randomizing the S-phase ordering of cells (right, **Methods**). The sigmoid fits are shown in red. The blue lines indicate the T-widths, defined as the time required for the replicated fraction to increase from 25% to 75%. **e)** Receiver operating characteristic curves of the trained eXtreme Gradient Boosting (XGB) models in RepTile’s machine-learning (ML) module evaluated using held-out G1-and G2-phase data and averaged across chromosomes (**Methods**). The true positive rate (TPR, y-axis) is plotted against the false positive rate (FPR, x-axis) at each classification threshold (**Methods**). The dashed line indicates random classification. The average area under the curve (AUC) is shown. **f-g)** Single-cell replication probability matrix for chromosome 6 predicted from RepTile (**f**) at 25kb resolution and measured by single-cell Repli-seq at 50 kb resolution (**g**) ^39^. Rows correspond to cells ordered by S-phase progression, and columns correspond to genomic loci. The locus-specific averages are shown on top of each matrix. **h)** Comparison of the percentage of replicated DNA inferred from each cell by RepTile’s ML module (y-axis) and RepTile’s statistical module (x-axis). The Pearson correlation is indicated.

We next inferred, for each S-phase cell, the fraction of loci that had replicated and used this value to order cells along S-phase progression ^39^ (**Figure 3b, Methods**). This inferred progression correlated strongly with nuclear volume (Spearman r_S_ = 0.82, p-value = 9.9e-123, **Figure 3c**), consistent with nuclear growth during S phase. This ordering also accurately recovered replication-timing heterogeneity, measured as T-width in independent Repli-seq experiments ^39,40^ (**Figure 3d** left and middle panels, **Methods**), whereas random ordering produced unrealistically broad T-width (**Figure 3d** right panel, **Methods**).

To assess the robustness and generalizability of RepTile, we applied it to an independent dataset in E14 mESC with substantially lower genomic coverage comprising 2,460 loci sampled at intervals of approximately 1Mb ^19^. Despite this sparse coverage, RepTile accurately inferred RT (Spearman *r_S_* = 0.75, p-value < 1e-300), S-phase progression and T-width (**Extended Figure 4a-e, Methods**).

### Assigning replication states to allele-resolved loci in single cells

We then applied RepTile’s machine-learning module (RepTile-ML) to infer the probability that each spatially resolved allelic locus copy in each individual cell had replicated (**Methods**). Chromosome-specific classifiers were trained on G1 and G2 cells, where replication states are known (unreplicated in G1 and replicated in G2) (**Methods**), and evaluated on a held-out test set. RepTile-ML showed good discrimination between replicated states (receiver operating characteristic area under the curve (AUC) = 0.83, **Figure 3e**, **Methods**). Importantly, this performance remained stable across loci stratified by RT, axial position within the coordinate frame (z-axis) or subnuclear location, indicating no detectable genomic or spatial biases (**Extended Figures 5abc, Methods**).

Applying RepTile-ML to S-phase cells generated a cell-by-locus matrix of imputed replication probabilities that recapitulated characteristic patterns observed by single-cell Repli-seq experiments ^39^ (**Figure 3fg**, **Methods**). Mean locus replication probabilities correlated strongly with replicated DNA fractions measured by single-cell Repli-seq (Pearson *r* = 0.87, p-value < 1e-300) (**Figure 3fg**, upper panels). Thresholding the imputed replication probabilities enabled binary replication state assignments, for example of 70% of loci at an estimated accuracy of 82% (**Extended Figure 5d, Methods**). These assignments also reproduced the RT measured by the ensemble Repli-seq experiment ^38^ (Pearson *r* = 0.88, p-value < 1e-300, **Extended Figure 5e**) and agreed closely with the RT (Pearson *r* = 0.93, p-value < 1e-300) and S-phase progression (Pearson *r* = 0.97, p-value < 1e-300) inferred by RepTile’s statistical module (**Figure 3h)**.

### Nuclear localization and its cell-to-cell variability correlate with replication timing

A key advantage of RepTile is that it relates replication dynamics directly to 3D genome organization in the same cells. For example, in a representative early-S cell, replicated loci cluster around nuclear speckles, supporting their proposed role as hubs for early-replicating DNA ^27,41^ (**Figure 4a**, left panel). As these cells progress into mid and late S, replicated loci accumulate more broadly throughout the nuclear interior away from speckles, while the remaining unreplicated loci become increasingly confined to the nuclear periphery, consistent with longstanding cytological observations ^42–44^ (**Figure 4a**, mid and right panels).

**Figure 4.**
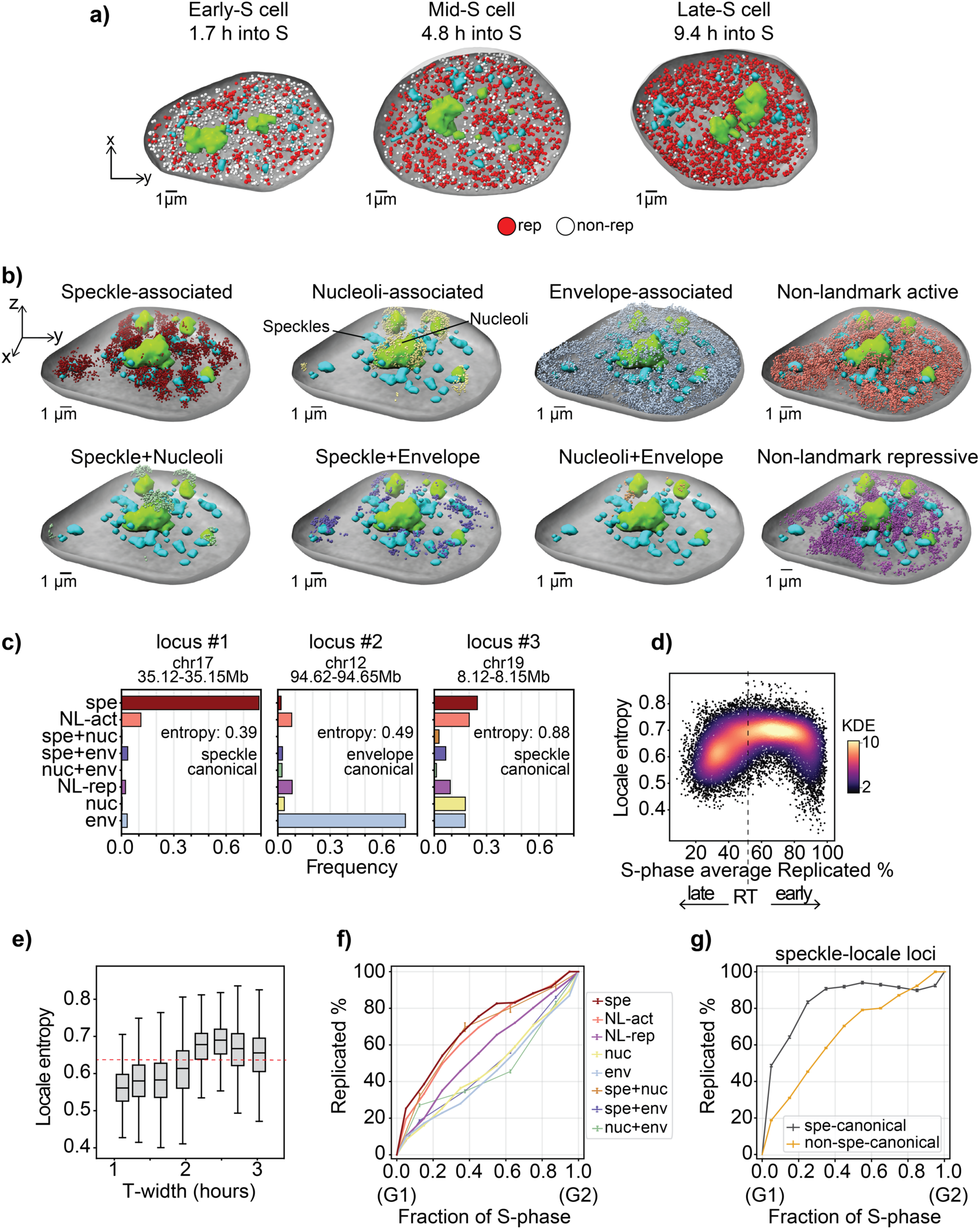
Spatially-defined sub-nuclear locales partition the genome by replication timing. **a)** 3D visualizations of three representative early-, mid-and late-S cells, showing replicated (red) and not-replicated (white) loci, together with speckles, nucleoli and the nuclear envelope. The cells are sectioned in the middle to allow internal visualization. **b)** Exploded view of a representative cell showing loci assigned to each nuclear locale separately, together with the nuclear envelope, speckles and nucleoli. **c)** Nuclear locale occupancy distributions for three representative loci, quantifying the fraction of allele instances assigned to each locale. The locale entropy, defined as the entropy value calculated from each distribution, is indicated together with the locus-specific canonical locale, defined as the most frequently occupied locale (**Methods**). **d)** Scatter-plot of the locus-specific locale entropy against locus-specific replication timing (RT) inferred from RepTile’s statistical module. Each dot, representing a 25kb region, is colored according to 2D Gaussian kernel density. **e)** Box-plots representing the locale entropy distributions of loci stratified by their T-width measured by high-resolution Repli-seq ^40^. The dashed red line indicates the genome-wide average locale entropy. **f)** Percentage of replicated DNA inferred from RepTile’s statistical module for loci assigned to each nuclear locale, calculated across cells grouped into ten consecutive S-phase windows (n=10 windows for: speckle, non-landmark active, non-landmark repressive, nucleoli, nuclear envelope locales; n=4 windows for mixed locales: speckle+nucleoli, speckle+envelope, nucleoli+envelope locales, **Methods**). The G1 and G2 fractions are reported as boundaries. **g)** As in **f**, but restricted to loci occupying the speckle locale and separated according to whether speckle was their canonical locale (black) or speckle was their non-canonical locale (orange).

To examine how replication relates to sub-nuclear location, we partitioned loci in each cell into quantiles based on their nearest distance to the nuclear envelope and the local SF3A66 (speckle) and Fibrillarin (nucleoli) fluorescence intensities sampled at each locus position, and estimated the fraction of replicated loci in each quantile (**Extended Figure 3f**, **Methods**). Replicated fractions increased with speckle marker intensity and were highest in the highest-intensity quantile (**Extended Figure 3f**, left panel). In contrast, replicated fractions peaked at intermediate nuclear envelope distances and intermediate nucleolar marker intensities, with the lowest replication fractions observed nearest to the nuclear envelope (**Extended Figure 3f**, middle and right panels). Because loci far from one landmark may simultaneously be close to another, intermediate distance effects cannot be uniquely attributed to a single landmark.

This motivated us to assign each locus in each cell to a ‘nuclear locale’ ^19,41,45,46^ based jointly on its relative positions to all three nuclear landmarks and its functional chromatin state. In each cell, we ranked loci by their SF3A66 (speckle) and Fibrillarin (nucleolus) fluorescence signals sampled at their positions and by their distance to the nuclear envelope. Then, we assigned locale associations using landmark-specific thresholds calibrated against independent TSA-seq data ^27^ (**Methods**). Loci not assigned to any landmark were further classified into active or repressive locales based on proximity to imaged active or repressive chromatin markers (**Methods**). Thus, each locus in each cell was assigned to one of eight nuclear locales: speckle-associated, nucleoli-associated, nuclear envelope-associated, joint speckle-nucleoli associated, speckle-nuclear envelope associated, and nucleoli-nuclear envelope associated, as well as non-landmark-active or non-landmark-repressive (**Figure 4b**, **Methods**).

Because any given locus can occupy different locales across cells, and even the two alleles of the same locus can occupy different locales in the same cell, we characterize each locus by a ‘nuclear locale occupancy profile’, defined as the fraction of its instances assigned to each locale across the cell population (**Figure 4c**). Some loci showed a strong preference for a single locale (**Figure 4c**, left and middle panels), while others were distributed more broadly across multiple locales (**Figure 4c**, right panel). To quantify this heterogeneity, we calculated for each locus the entropy of its nuclear occupancy profile (**Methods**): low entropy indicates strong preference for a particular locale (**Figure 4c**, left panel), while high entropy indicates a more uniform distribution across all locales (**Figure 4c**, right panel). Additionally, for each locus we identified its ‘canonical locale’, defined as the most frequently visited locale among all its allele instances (**Figure 4c**, **Methods**).

We found that the earliest-and latest-replicating loci had the lowest entropy, while mid-replicating loci had higher entropy values (**Figure 4d**). We also found that loci with greater variability in replication timing, as measured by locus-specific T-width in high-resolution Repli-seq experiments ^40^, showed higher locale entropy (Welch t-test t = 79.4, d.f. = 18,900, p-value < 1e-300, **Figure 4e**, **Methods**). Thus, the earliest-and latest-replicating loci not only occupied a more specialized nuclear environment than mid-replicating loci but also showed lower replication timing variability. This suggests that stable positioning in defined nuclear microenvironments is associated with more precise replication timing, while broader positional plasticity accompanies stochasticity in locus firing.

To study how replication timing varies across nuclear locales, we ordered S-phase cells by their replication progression and divided them into 10 consecutive intervals. Within each interval, we determined the fraction of loci in each locale that had replicated, generating locale-specific cumulative replication curves (**Figure 4f, Methods**). These curves allowed us to compare the relative replication timing across nuclear locales and determine how broadly replication was distributed across S phase (**Figure 4f**). Speckle-associated loci showed the highest replicated fractions, consistent with early replication. Loci in the non-landmark active locale also replicated early, but significantly later than speckle-associated loci (z-test z = 16.0, p-value = 2.9e-57). For example, in the earliest S-phase interval, 25.2% of speckle-associated loci had replicated, compared with 19.4% of non-landmark-active loci. Speckle-nucleoli loci also showed relatively high replicated fractions, although lower than speckle-associated loci (z = 4.8, p-value = 1.4e-6). In contrast, speckle-envelope, nucleolar and nuclear-envelope locales showed substantially lower replicated fractions, whereas loci in the non-landmark repressive locale showed an intermediate replication behavior (**Figure 4f**). Among repressive locales, nuclear envelope-associated loci consistently showed the lowest replicated fractions across all intervals (all z ≥ 5.8, p-value < 6.5e-9) (**Figure 4f**).

Our results are consistent with extensive prior findings from population-level analyses linking speckle proximity and active chromatin to early replication, and lamina or nucleolar proximity and inactive chromatin to late replication ^1,42,43^. However, these previous studies relied either on correlations between independent bulk genomics experiments, such as Repli-seq and TSA-seq ^27,41^, or on classical cytological imaging measurements such as BrdU staining, which lack any locus-specific information ^44,47^. In contrast, our method provides a direct, single-cell and locus-resolved analysis from a single imaging experiment. In this context, our results can be viewed as the first direct assessment of the prior indirect genomics and cytological approaches.

Interestingly, we also observed that approximately 80% of speckle-associated loci had replicated by the midpoint of S phase, whereas the replication times of the remaining 20% spanned the entire second half of S phase, indicating that speckle association does not imply uniformly early replication for every locus (**Figure 4f**). To examine the basis of this heterogeneity, we separated all speckle-associated loci in each S-phase interval into two groups: those that had the speckle environment as their preferred canonical location across the population of cells and those that showed non-speckle-canonical locales, and thus were more frequently associated with other locales but occupied the speckle locale in the cell being analyzed. We then determined the fraction of replicated loci separately for both groups across all S-phase intervals, generating two cumulative replication curves (**Figure 4g**). We found that canonical speckle loci showed consistently higher replicated fractions during early and mid S-phase than non-canonical speckle loci (z ≥ 16.9, p-values < 1.7e-64 for all intervals except the final three intervals, when both groups were highly replicated). This indicates that the relationship between speckle association and early replication timing is not uniform across loci but rather depends on the locus-specific preferences for that locale. We examine this dependence systematically in the next section.

### Nuclear positioning influences single-cell replication timing in a locus-specific manner

So far, we have analyzed replication probabilities across nuclear locales irrespective of locus identity. We next asked whether the replication differences observed across locales reflect the intrinsic properties of loci that preferentially occupy those locales, or whether the nuclear locale itself is a major driver of replication probability. In the latter case, the same locus would change its replication behavior with its nuclear location in single cells. We therefore compared loci that occupied different locales across single cells and asked whether their replication probability changed with their location. For example, **Figure 5a** shows a locus (chr5:138.0-138.2Mb) in a cell where it was replicated when positioned near a speckle but remained unreplicated in another cell at a similar stage of S-phase when positioned near the nuclear envelope. To quantify this effect across the full cell population, we compared the locale occupancy profile of its replicated instances with that of its unreplicated instances. The two profiles differed significantly (Chi^2^ = 147.3, p-value = 7.7e-31, d.f. = 4, **Figure 5b**, left panel): 93% of replicated instances occupied the speckle or non-landmark active locales, whereas 57% of unreplicated instances were located at the nuclear envelope, indicating that replication probability for this locus depends on the nuclear locale it occupies. However, for other loci the difference between replicated and unreplicated locale occupancy profiles was only marginal, indicating that the relationship between nuclear locale and replication probability is not uniform across loci. For example, locus chr17:70.2-70.4Mb or locus chr2:24.2-24.4Mb showed no significant differences in the locale occupancy profiles of their replicated and unreplicated instances (Chi^2^ = 3.2, p-value = 0.52, d.f. = 4, and Chi^2^ = 1.4, p-value = 0.84, d.f. = 4, **Figure 5b**, mid and right panels). These locus-specific differences in locale dependence indicate that nuclear position is not a universal determinant of replication timing but instead acts selectively on specific classes of loci.

**Figure 5.**
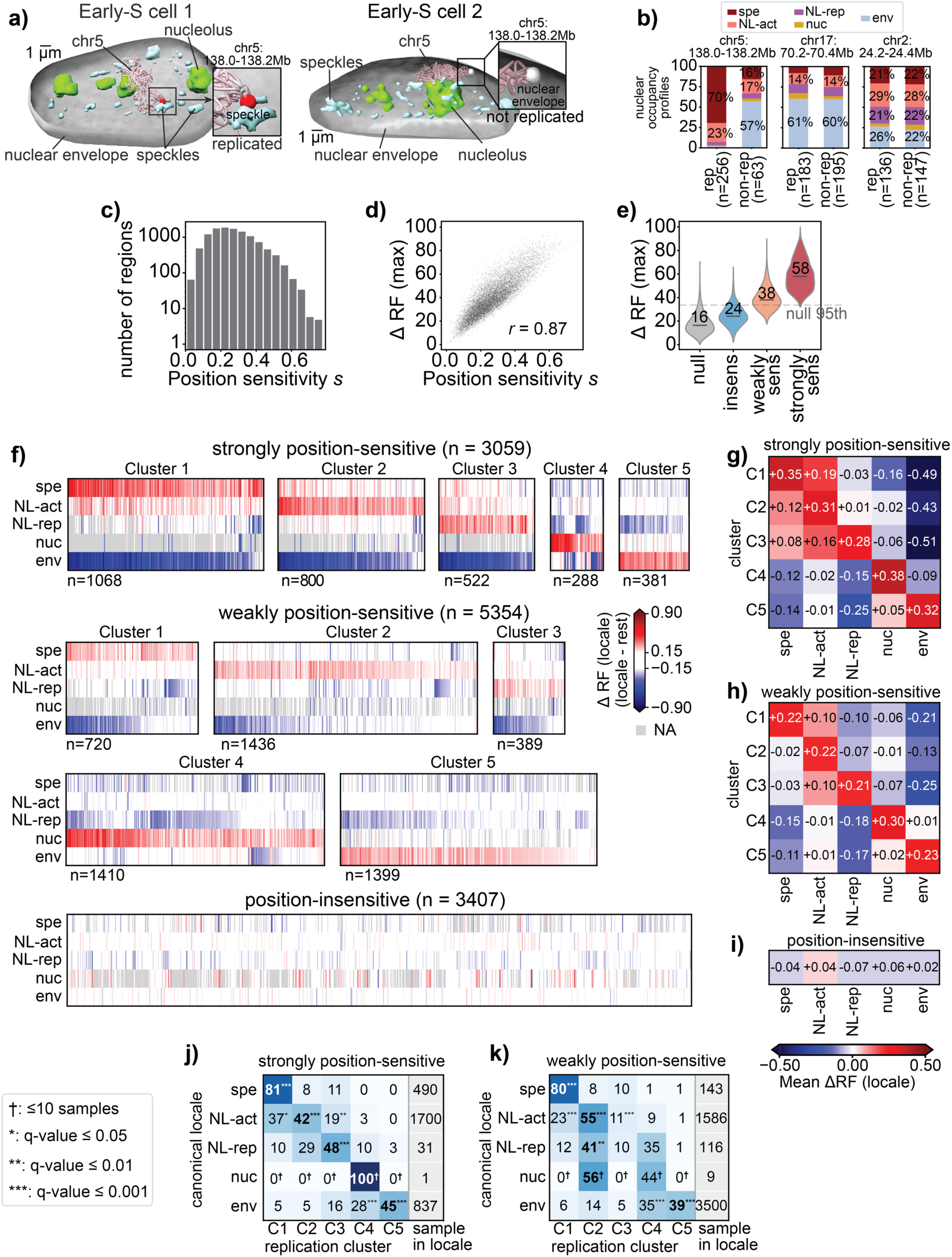
Locus-specific sub-nuclear locales modulate the replication propensity of ‘position-sensitive’ regions. **a)** 3D visualizations of two representative early-S-phase cells at a similar S-phase stage showing the nuclear envelope (grey), speckles (cyan) and nucleoli (green), one copy of chromosome 5 (pink) and a highlighted locus (ch5:138.0-138.2Mb, red/white beads). RepTile predicted the locus was replicated in the cell on the left (left) and unreplicated in the cell on the right (right). **b)** Bar-plots showing the distributions of the nuclear occupancy profiles of three representative loci. For each locus the nuclear occupancy profiles are shown separately for replicated and unreplicated allele instances in S-phase cells. Sample sizes are indicated for each locale occupancy profile. The locus on the left is the locus shown in **a**. **c)** Genome-wide distribution of position-sensitivity values for loci at 200kb resolution (**Methods**). **d)** Genome-wide scatter plot relating locus-specific position sensitivity to the maximum difference in replicated fraction between allele instances of the same locus occupying any two nuclear locales ΔRF(max), for each 200kb locus across S-phase cells (**Methods**). The Pearson correlation is indicated. **e)** Distribution of ΔRF (max) for strongly position-sensitive, weakly position-sensitive and position-insensitive loci, together with a null distribution obtained by randomizing locale assignments (**Methods**). Numbers indicate median values, and the dashed line corresponds to the 95^th^ percentile of the null distribution. **f)** Heatmaps showing the locale-associated replication shift for all position-sensitive and position-insensitive loci. Each column represents a 200kb locus and each row one of the five primary locales (speckle, non-landmark active, non-landmark repressive, nucleoli, envelope), such that each locus is represented by five Δ*RF_i_* (locale) values. Δ*RF_i_* (locale) is defined as the difference between the fraction of replicated instances of locus *i* when it occupied a given locale and the fraction replicated when it occupied any other locale (**Methods**). Position-sensitive loci were clustered into groups based on their five-value Δ*RF_i_* (locale) vectors, and entries are colored based on Δ*RF_i_* (locale) (**Methods**). The five clusters are shown separately for strongly (top) and weakly (middle) position-sensitive loci. Position-insensitive loci were not clustered because they showed only minor locale-associated differences in replication. **g-h-i)** Mean ΔRF(locale) across loci in each cluster and for each primary locale, shown separately for strongly position-sensitive (**g**), weakly position-sensitive (**h**) and position-insensitive (**i**) loci. Position-insensitive loci are represented by a single row because they were not clustered. **j-k)** Matrices showing the percentage of loci with each canonical locale (rows) assigned to each of the five replication-locale clusters (columns), separately for strongly (**j**) and weakly (**k**) position-sensitive loci. Rows are normalized to 100%, and the sixth column indicates the total number of loci for each canonical locale. Bold values indicate the largest percentage in each row. Stars indicate Benjamini–Hochberg-corrected q-values from one-sided permutation tests for over-representation relative to a null model of independence between canonical locale and cluster (*q < 0.05, **q < 0.01, ***q < 0.001, **Methods**). Daggers (†) indicate percentages calculated from fewer than ten nucleolus-canonical loci.

To quantify this effect systematically genome-wide, we implemented a statistical Analysis of Variance (ANOVA) approach (**Methods**) based on multilinear regression to estimate for each locus the fraction of replication state variation that can be attributed to differences in nuclear locale occupancy across single cells. We quantified this fraction for each locus as the position-sensitivity score, *s* (**Methods**). A locus with high positional sensitivity *s* showed strong locale-dependent replication, while loci with low *s* replicated at similar rates regardless of their nuclear environment. Across 11,820 genomic regions at 200kb resolution, *s* spanned a broad spectrum, from values close to zero to as high as 0.78 (**Figure 5c**, **Methods**).

To confirm *s* as a measure of replication position-sensitivity, we calculated, for each locus, the maximum difference in the replicated fraction (Δ*RF_i_* (max)) between any two nuclear locales. Specifically, for each locus all its instances were grouped by the locale they occupied, and for each locale the replicated fraction was determined across all S-phase cells. We then defined Δ*RF_i_* (max) as the maximum difference in replicated fraction between any two nuclear locales for locus *i* (**Methods**). As expected, *s* correlated strongly with Δ*RF_i_* (max) (Pearson *r* = 0.87, p-value < 1e-300, **Figure 5d**), confirming that the relationship between nuclear organization and replication timing is locus-specific rather than uniform, allowing distinct classes of loci to be distinguished by their degree of positional dependence.

### Genomic regions group by locale-dependent patterns of single-cell replication timing

Next, we divided genomic loci into three classes based on their position-sensitivity *s*. Loci with *s* < 0.19, whose observed sensitivity did not exceed the permutation-based null expectation, were classified as ‘position-insensitive’ (n = 3,407, 28.8%; Benjamini-Hochberg FDR > 0.05; **Figure 5e** and **Methods**). The remaining loci were classified as position-sensitive, and further divided into ‘strongly’ (*s* > 0.34; n = 3,059, 25.9%) and ‘weakly’ (0.19 < *s* < 0.34; n = 5,354, 45.3%) position-sensitive groups, thereby separating high-and low-sensitivity groups for categorical analyses (**Methods**).

While *s* quantifies how strongly replication depends on a locus’s nuclear location, it does not reveal which nuclear locales advance or delay replication. We therefore investigated which specific locales were associated with the earliest or latest replication for position-sensitive loci. For each locus and locale, we calculated Δ*RF_i_* (locale) as the difference between the fraction of locus instances that were replicated when the locus occupied that locale and the fraction replicated when it occupied any other locale. Δ*RF_i_* (locale) is therefore a measure of whether replication for a specific locus is advanced or delayed in that nuclear locale environment compared to all other locales (**Methods**). Thus, each locus (*i*) was represented by a vector of five locale-specific Δ*RF_i_* (locale) values, one for each primary locale, which together define its locale-associated replication shift. Clustering position-sensitive loci based on Δ*RF* vectors identified five groups with distinct patterns of locale-dependent replication (**Figure 5f**, **Extended Figure 6a, Methods**).

The five clusters differed primarily in the nuclear locale associated with maximal replication advancement. Cluster 1 loci (C1) showed the highest fraction of replication, and thus the largest positive replication shift, when positioned near speckles, with mean ΔRF_C1_(speckle) values of 35% and 22% for strongly and weakly position-sensitive loci, respectively (**Figure 5f**, top row, **Figure 5gh**). In contrast, the same loci were markedly delayed in replication at the nuclear envelope, with mean ΔRF_C1_(envelope) values of -49% and -21%. Thus, loci that replicate among the earliest near speckles could become among the latest replicating when positioned at the nuclear envelope (**Figure 5fgh**). These loci also showed substantially lower mean ΔRF_C1_ values in non-landmark active, non-landmark repressive, and nucleolar locales than in the speckle-associated locale (**Figure 5fgh**).

Surprisingly, speckle association was not the locale of maximal replication advancement for most position-sensitive loci. Cluster 2 loci (C2) replicated earliest in the non-landmark-active locale, with mean ΔRF_C2_(non-landmark active) values of 31% and 22% for strongly and weakly position-sensitive loci, respectively, compared with 12% and -2% near speckles (ΔRF_C2_(speckle)) (**Figure 5fgh**). Nuclear envelope association reduced the replicated fraction further for these loci with mean ΔRF_C2_(envelope) values of -43% and -13% for strongly and weakly position-sensitive loci. Cluster 3 loci (C3) showed maximal replication advancement in the non-landmark-repressive locale with mean ΔRF_C3_(non-landmark repressive) values of 28% and 21%, for strongly and weakly position-sensitive loci, demonstrating that a repressive environment could favor earlier replication over speckle-associated or other active environments (**Figure 5fgh**). Despite their earlier replication in a repressive environment, cluster 3 loci showed also a substantial negative replication shift at the nuclear envelope, particularly among strongly position-sensitive loci (ΔRF_C3_(envelope) = -51% and -25% for strongly and weakly position-sensitive loci). Cluster 4 loci (C4) replicated earliest in the nucleolar locale (ΔRF_C4_(nucleoli) = +38% and 30% for strongly and weakly position-sensitive loci) and were delayed in all other locales (**Figure 5fgh**). Finally, cluster 5 loci (C5) showed the opposite behavior of cluster 1: replication was earliest at the nuclear envelope (ΔRF_C5_(env) = +32% for strongly position-sensitive loci), but was substantially delayed near speckles and in the non-landmark-repressive locale (ΔRF_C5_(speckle) = -14%, ΔRF_C5_(non-landmark repressive) = -25%). Weakly position-sensitive loci showed similar locale preferences, although with smaller replication shifts. (see **Extended Figure 6b** for a combined analysis of all position-sensitive loci).

As expected, position-insensitive loci showed little evidence of systematic shifts in replication timing across nuclear locales, with only marginal differences in ΔRF (locale) across locales (**Figure 5f**, lowest panel, and **Figure 5i**). Even speckle or nuclear envelope associations, which at the population level are linked to the earliest and latest replication timing, had little or no detectable effect on replication timing.

Together, these cluster-specific patterns show that nuclear locales do not impose a uniform effect on replication timing. Instead, the same locale can advance replication for one class of loci while delaying replication for another, revealing a locus-specific relationship between nuclear position and replication timing.

### Canonical nuclear positioning confers a replication-timing advantage for position-sensitive loci

The distinct cluster-specific patterns raised the question of why different loci replicated earliest in different nuclear environments. We therefore asked whether each locus’s canonical locale, i.e. the locale it occupied most frequently across cells, corresponded to its replication-favorable locale (RFL), defined as the locale with the largest positive replication shift of the cluster it was assigned to. Among strongly position-sensitive loci, the RFL matched the canonical locale significantly more often than expected by chance (49.1% vs a null of 23.7%, observed/expected O/E = 2.07, permutation test p-value < 1e-4, **Methods**). This relationship increased with the “canonical strength”, defined as the ratio between the frequencies of the canonical and second-most frequent locales. Among loci with a canonical strength of at least 2, 63.3% had matching canonical and replication favorable locales, compared with 26.0% expected under the null model (O/E = 2.44, p-value < 1e-4, **Methods**). This correspondence was present across all canonical locales with sufficient representation. For loci within each canonical locale, the largest fraction of loci belonged to the cluster whose RFL matched that canonical locale (**Figure 5j**). Specifically, 81% of speckle-canonical loci belonged to C1 (O/E = 2.32), which had speckle as its RFL; 42% of non-landmark active loci to C2 (O/E = 1.59), which had non-landmark-active as its RFL; 48% of non-landmark repressive loci belonged to C3 (O/E = 2.84), which had non-landmark-repressive as its RFL; and 45% of envelope-canonical loci belonged to C5 (O/E = 3.64), which had the nuclear envelope as its RFL. All associations were highly significant under permutation tests (Benjamini-Hochberg q-value = 5e-4). Nucleolus-canonical loci were too scarce for statistical testing (n = 1) and were excluded.

We also observed weaker secondary preferences for some locales. For example, non-landmark active-canonical loci were additionally represented in C1 (RFL = speckle, 37%, O/E = 1.05, q-value = 0.05) and C3 (RFL = non-landmark repressive, 19%, O/E = 1.10, q-value = 0.01), whereas envelope-canonical loci showed a secondary presence in C4 (RFL = nucleoli, 28%, O/E = 2.98, q-value = 5e-4). In each case, however, the cluster whose RFL matched the canonical locale remained the strongest association. Weakly position-sensitive loci showed a similar pattern (**Figure 5k**, **Extended Figure 6c** shows all position-sensitive loci combined), except that non-landmark repressive-canonical loci were more frequently assigned to cluster C2. Thus, although loci in different clusters showed replication timing advantage in different nuclear environments, they shared a common organizing principle: loci tended to replicate earliest in their preferred canonical spatial environment.

### Within-cell allele comparisons indicate that variation of cell-intrinsic factors does not explain the replication advantage of replication-favorable locales

Our analysis so far compared the replication state of the same locus across a population of cells, and therefore often across different cellular contexts. A key concern is that the observed association between nuclear locale and replication state could be confounded by cell-to-cell variation in intrinsic properties of the cell, such as nuclear volume, global replication progression, or factor abundance during S-phase, rather than reflecting a direct effect of the locale nuclear environment. To control for these sources of variation, we identified specific cells in which the two alleles of a given locus were in discordant replication states, with one allele replicated and the other unreplicated (**Figure 6a**, **Methods**). We refer to these as discordant-allele cells for that locus. Comparing alleles within the same cell ensures that both copies experienced the same global cellular context.

**Figure 6.**
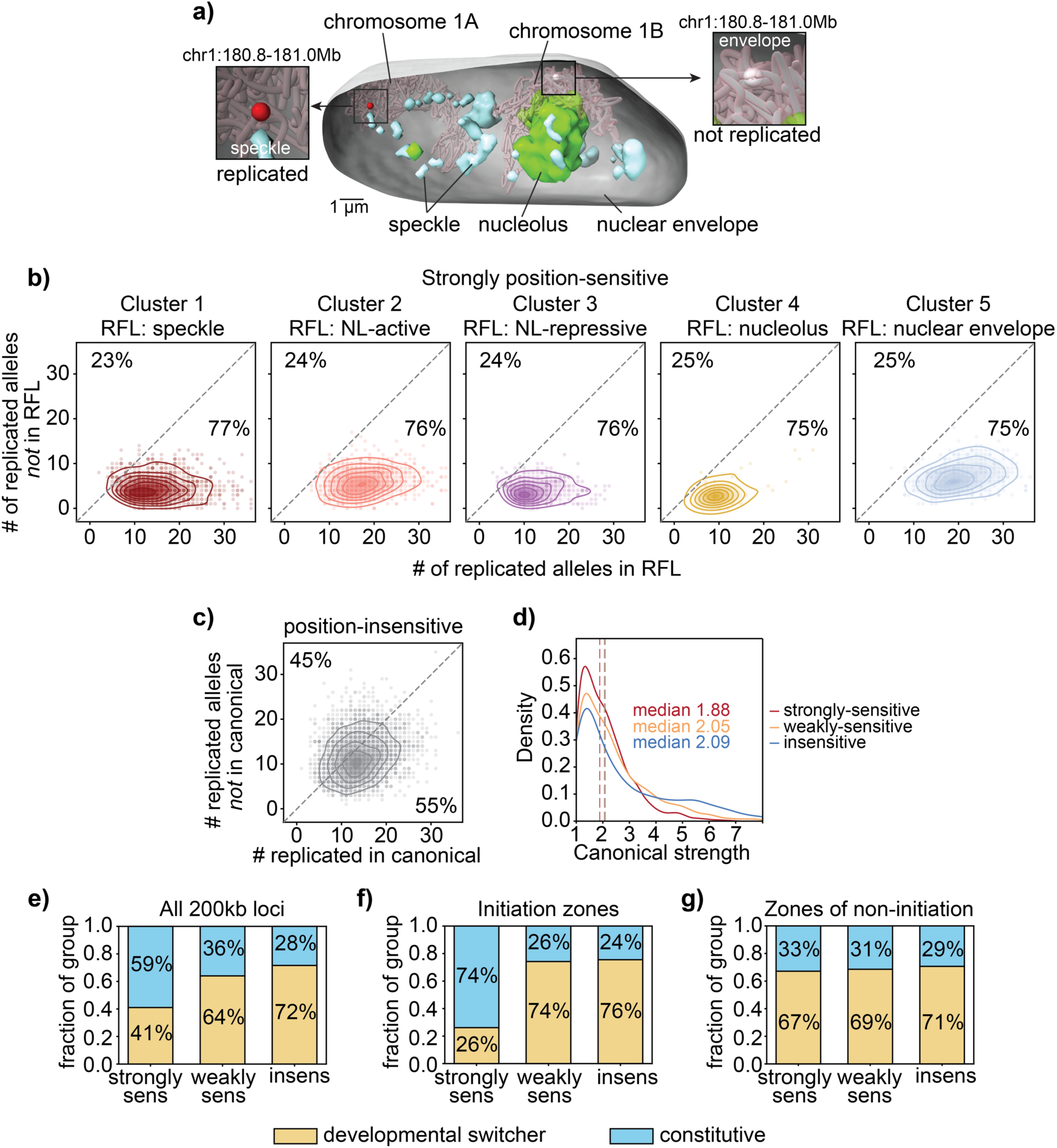
Position-sensitive initiation zones preferentially overlap with constitutive replication timing domains, while position-insensitive zones overlap with developmental switchers. **a)** 3D visualization of a representative early-S cell showing the nuclear envelope (grey), speckles (cyan) and nucleoli (green), together with both copies of chromosome 1 (pink). Red and white beads mark the two alleles of a locus (chr1:180.8-181.0Mb), colored according to whether RepTile predicted them to be replicated (red) or not-replicated (white), respectively. **b)** Locus-specific scatter and density contour plots for strongly position-sensitive loci, showing the number of discordant-allele cells, i.e. those in which one allele was replicated and the other unreplicated and in which the replicated allele occupied the locus’s replication-favorable locale (RFL, x-axis, **Methods**) or another locale (y-axis). The dashed line indicates equal frequencies. Percentages below and above the plots indicate the fractions of allele-discordant cells in which the replicated allele occupied the RFL or another locale, respectively. Loci are grouped by replication–locale cluster (**Methods**). **c)** As in **b**, for position-insensitive loci. However, because these loci do not have a well-defined RFL, their canonical locale was used instead. **d)** Distribution of canonical strength, defined as the ratio between the occupancy frequencies of the canonical locale and the second most-frequent locale (**Methods**), for strongly position-sensitive (red), weakly position-sensitive (orange) and position-insensitive (blue) loci (**Methods**). Dashed vertical lines indicate medians. **e-f-g)** Fractions of strongly position-sensitive, weakly position-sensitive and position-insensitive loci overlapping domains that replicate constitutively early or late across 31 mouse cell types (cyan) or switch replication timing during development (yellow) ^3,48^ (**Methods**). Fractions are shown genome-wide (**e**), within conserved mESC E14 initiation zones (**f**), and within conserved zones of non-initiation (**g**, **Methods**).

We then quantified, among discordant-allele cells, how often the replicated allele occupied the replication-favorable locale of the cluster to which that locus belonged (**Figure 6b, Extended Figure 6d**). Consistent with the analysis across all locus instances and cells, the replicated allele of strongly position-sensitive loci occupied the cluster-specific replication-favorable locale significantly more often than expected by chance, exceeding the 50% null (**Figure 6b**). For instance, among cluster 1 loci, the replicated allele occupied the replication-favorable locale in a median of 77% of discordant-allele cells (one-sample Wilcoxon signed-rank vs null p-value = 6.7e-152), compared with 76% for cluster 2 loci (Wilcoxon p-value = 1.6e-130), with the remaining clusters at either 75% or 76% (**Figure 6b**). In contrast, position-insensitive loci showed only weak tendencies of their replicated alleles to be in a specific locale, as for instance the loci’s canonical locale (55% vs 45%) (**Figure 6c**, **Methods**). These within-cell allele comparisons indicate that the locale-replication association of position-sensitive loci cannot be explained by cell-to-cell variation of cell intrinsic factors.

### Position-insensitive loci are enriched for domains with developmentally regulated replication timing

We next asked what distinguishes position-insensitive loci, whose replication timing is largely independent of their nuclear position, from strongly position-sensitive loci, whose replication is tightly coupled to subnuclear location. Because canonical positioning conferred a replication-timing advantage for position-sensitive loci, we first tested whether position-insensitive loci simply showed weaker canonical positioning and thus lower canonical strength. This was not the case: the median canonical strength was higher for position-insensitive loci (2.20) than for position-sensitive (strongly: 1.82, weakly: 2.05; strongly vs insensitive Mann-Whitney p-value = 7.1e-57). This difference was driven in part by a more pronounced upper tail of loci with high canonical strength in the position-insensitive group (**Figure 6d**). Thus, position-insensitive loci can retain strong canonical locale preferences while their replication timing remains largely independent of their nuclear position.

We next compared the genomic and epigenetic features of position-sensitive and position-insensitive loci. Position-sensitivity showed modest positive correlations with active chromatin signatures (Spearman correlation ρ = 0.31–0.37, across several active chromatin markers, **Extended Figure 6e-top**, **Methods**). However, these associations were largely attenuated after controlling for the canonical locale by computing partial correlations (partial Spearman correlations ρ = 0.01-0.07, **Extended Figure 6e-bottom**, **Methods**), indicating that the positive correlation between position-sensitivity and active markers was mostly driven by canonical locale imbalances across high and low position-sensitivity groups.

No tested individual epigenetic feature clearly distinguished position-sensitive from position-insensitive loci, indicating that position sensitivity is not determined by a single genomic or chromatin marker. However, we observed a striking association with replication-timing plasticity during development. Position-insensitive loci were strongly enriched for domains that switch between early and late replication across 31 mouse cell types ^3,48^, while strongly position-sensitive loci preferentially overlapped domains that remain constitutively early or late replicating (strongly position-sensitive vs position-insensitive odds ratio = 4.30, p-value = 3.1e-137, **Figure 6e**, **Methods**). This association persisted after stratifying loci by replication timing (Mantel-Haenszel odds-ratio: 2.53, p-value: 1.4e-35, **Methods**) or by canonical location (Mantel-Haenszel odds-ratio: 2.29, p-value: 2.0e-29), indicating that the enrichment cannot be explained solely by replication timing or nuclear position.

Because replication timing is ultimately established through the temporal regulation of origin firing, we next asked whether the association between position sensitivity and developmental replication-timing plasticity was specifically linked to initiation zones, as would be expected if the regulation is at the level of replication initiation. In mammals, the most efficient replication origins occur within broad initiation zones (IZs), ∼45kb regions that contain clusters of initiation events at elevated frequency ^49,50^. We therefore curated a set of high confidence IZs in E14 mESC cells by identifying peaks in log_2_(early/late) Repli-seq signal ^38^ and intersecting these candidates with high resolution Repli-seq IZs from hybrid (castaneus x musculus) F121-9 mESC, and allele-resolved *mus musculus* ^40^ (**Extended Figure 7ab**, **Methods**). Convincingly, this intersection yielded 895 conserved IZs (42% of IZs across all datasets). We also defined conserved non-IZ regions to test whether the association between position sensitivity and developmental replication-timing class was specific to origin-containing regions (**Methods**).

Strikingly, the enrichment of strongly position-sensitive regions in constitutively replicating domains was driven almost entirely by IZs. Among strongly position-sensitive IZs, 75% overlapped constitutively early or constitutively late domains (**Figure 6f**). In contrast, weakly position-sensitive and position-insensitive IZs were predominantly located in developmentally switching domains (66% and 85% overlap, respectively, **Figure 6f**). Outside IZs this distinction was lost: strongly position-sensitive, weakly position-sensitive and position-insensitive non-IZs all showed similar enrichment for developmentally switching domains, at approximately 70% (**Figure 6g**). Thus, the association between strong positional control and constitutive replication timing is not a general property of chromosomal regions but likely reflects the regulation of replication initiation.

Developmentally regulated replication-timing domains often re-localize within the nucleus during differentiation, in coordination with changes in transcription and chromatin state, while constitutively early and constitutively late domains tend to retain their subnuclear positions across cell types ^51–54^. Developmentally switching domains may rely more strongly on intrinsic regulatory mechanisms that decouple their replication program from differentiation-driven structural rearrangements. By contrast, constitutive domains may be more permissive to positional control, because their nuclear environments are comparatively stable across cell types (see **Discussion**).

To search for direct evidence of intrinsic regulatory elements at position-insensitive loci, we examined three IZs for which early-replicating control elements (ERCEs) have been experimentally validated by CRISPR deletion ^38^. ERCEs are cis-regulatory regions whose deletion substantially delays the replication timing of their associated IZs, providing causal evidence for intrinsic replication-timing control. All three ERCE-validated loci, *Dppa2*, *Zfp42/Rex1*, and *Klf4*, were position-insensitive in our framework, with *s* values of 0.167, 0.168 and 0.177, respectively, all below the insensitivity threshold of 0.19. Although the small number of functionally validated ERCEs precludes statistical inference, this pattern is consistent with the hypothesis that position-insensitive IZs are more likely to harbor intrinsic cis-regulatory elements capable of controlling replication timing independently of nuclear locale. More broadly, since ERCEs, like transcriptional facilitators ^55,56^, are difficult to distinguish from other active regulatory features without perturbation, additional functional assays will be needed to determine whether intrinsic replication-timing control is a general property of position-insensitive IZs.

## Discussion

Here we introduce RepTile, a systematic framework for inferring locus-and chromosome-copy-resolved replication states directly from multiplexed DNA FISH and chromosome-tracing data ^9,10,13,16,19^. RepTile addresses a fundamental ambiguity in the analysis of proliferating cells: observed spot counts reflect both DNA copy number and substantial locus-and position-dependent variation in detection. By treating replication state as a latent variable and explicitly accounting for detection efficiency, background signal and overcounting, RepTile separates biological changes in copy number from technical variation in imaging and barcode decoding. Its statistical and machine-learning modules together infer cell-cycle stage, S-phase progression, replication timing, cell-to-cell timing heterogeneity and the replication states of individual spatially resolved locus copies within their native three-dimensional nuclear context.

Resolving replication state is important not only for studying DNA replication itself, but also for the quantitative interpretation of multiplexed imaging data. Replication changes the number, intensity and spatial extent of locus signals and can therefore bias chromosome-copy assignment, locus–locus distances, estimates of chromatin compaction and chromosome-territory volume, and associations between genome structure, nuclear position and transcription. RepTile provides a principled means of accounting for this latent copy-number variation rather than excluding S-phase cells or relying on heuristic identification of replication doublets. More broadly, its ability to infer cell-cycle progression directly from the imaging data enables structurally resolved nuclei to be ordered without prospective synchronization, allowing genome organization to be examined continuously across S phase. Because the framework operates on decoded locus-level measurements, it is not intrinsically limited to complete genome coverage in every cell and is applicable to datasets targeting whole genomes, individual chromosomes or selected chromosomal regions, provided that sufficient measurements are available for calibration and inference.

The close agreement of RepTile-derived replication timing with independent ensemble and single-cell Repli-seq measurements ^38–40^ demonstrates that replication information is recoverable from multiplexed imaging despite extensive detection variability. Importantly, direct use of uncorrected spot counts produced substantially poorer agreement with Repli-seq, showing that the inferred replication patterns cannot be obtained by simple spot counting alone. RepTile also recovered S-phase progression and replication-timing heterogeneity and assigned replication states to individual allelic copies with comparable performance across loci differing in replication timing and spatial position. These results establish multiplexed imaging as a source to study replication while providing spatial measurements that are unavailable from sequencing-based assays, including allele-specific three-dimensional position and proximity to nuclear structures.

By resolving the replication states of individual alleles within their native nuclear context, RepTile enables replication timing to be examined together with chromatin folding and subnuclear positioning in the same cell. Such joint, genome-scale measurements have not previously been possible and provide a direct means to assess how nuclear organization influences replication decisions at individual loci. This, in turn, provides insight into the relative contributions of local nuclear environment and locus-intrinsic regulatory programs to replication timing control.

Previous population-level and classical cytological imaging studies have associated speckle proximity and active chromatin with early replication, and lamina or nucleolar association with late replication ^1,27,41–44,47^. Consistent with these observations, speckle-associated loci showed the highest replicated fractions across S phase, whereas envelope-associated loci showed the lowest. However, our locus-resolved single-cell analysis revealed that these population-level associations conceal fundamentally different behaviors among genomic loci.

First, nuclear position does not exert a uniform influence on replication timing for all loci across the genome. Replication timing for some loci was strongly position sensitive and changed markedly with their subnuclear location, whereas other loci replicated at similar times irrespective of their subnuclear environments. Importantly, this distinction was not explained by the strength of a locus’s spatial preference: position-insensitive loci could have canonical positioning preferences as strong as, or stronger than, those of position-sensitive loci. More broadly, the coexistence of position-sensitive and position-insensitive loci underscores that occupancy of a particular nuclear environment is not inherently regulatory; rather, its functional consequences depend on the identity and regulatory state of the locus. Thus, population-level associations between nuclear position and replication timing can arise from loci that are highly responsive to nuclear position and loci whose replication timing is largely independent of it.

Second, not all position-sensitive loci replicated earliest at speckles or latest at the envelope, indicating that these environments do not universally support early or late replication. Only a subset of loci replicated earliest at speckles or latest at the lamina; some loci showed delayed replication when located at speckles or earliest replication at the lamina. We identified five groups of position-sensitive loci with distinct locale-dependent replication patterns, each replicating earliest in a specific nuclear locale. We conclude that the same nuclear environment can advance replication for one class of loci while delaying it for another. Consistent with our finding, pericentromeric constitutive heterochromatin has previously been shown to replicate earlier when tethered to the nuclear lamina ^57^.

Third, despite this diversity, position-sensitive loci shared a broader organizing principle: replication was generally most advanced in the locale that the locus preferentially occupied. This correspondence between canonical and replication-favorable locales suggests that position-sensitive genomic regions are functionally matched to their preferred nuclear environments rather than responding uniformly to a universal hierarchy of active and repressive compartments. Position-insensitive regions have equally preferable locales, but do not have a replication-favorable locale.

Fourth, these principles were the same between cells or between homologous alleles within the same cell, demonstrating that they were not dependent upon variation of cell intrinsic variables, such as nuclear size, factor abundance or S-phase progression.

Fifth, we found that strongly position-sensitive initiation zones were preferentially associated with domains that replicate constitutively at the same time across cell types, whereas position-insensitive initiation zones were enriched in developmentally regulated domains that switch replication timing during differentiation. This relationship between constitutive domains and position-sensitivity was largely absent outside initiation zones, indicating that it is specifically associated with the regulation of replication initiation rather than being a general property of constitutive and developmentally regulated chromatin. These findings suggest two broad, and potentially complementary, modes of replication control. Constitutive initiation zones may rely more strongly on developmentally stable nuclear environments that support their characteristic replication timing. By contrast, developmentally regulated initiation zones may possess stronger locus-intrinsic regulatory programs that allow their replication timing to remain buffered from their immediate nuclear position.

Such intrinsic regulation could be particularly important during differentiation, when developmentally regulated domains frequently change chromatin state, transcriptional activity and subnuclear position. Dependence on a single spatial environment could constrain their ability to switch replication timing as cell identity changes. Position-insensitive initiation zones may instead carry cis-regulatory information that allows their replication program to be reset independently of nuclear locale. Such mechanisms could explain how replication timing and nuclear position can be coordinately re-established during early G1 while remaining mechanistically separable at a subset of developmental domains. Consistent with this model, all three initiation zones containing experimentally validated early replication control elements (ERCEs) ^38^ were classified as position insensitive. The small number of functionally validated elements precludes a general conclusion, but the observation raises the possibility that ERCEs and related cis-regulatory elements confer positional independence. Systematic perturbation of candidate elements will be needed to determine whether locus-intrinsic control is a general feature of developmentally regulated initiation zones.

These findings refine the mechanistic interpretation of the timing decision point (TDP) ^3^, at which replication timing and nuclear organization are coordinately re-established in early G1. This temporal coincidence has long supported the idea that subnuclear position contributes to the establishment of replication timing. Our results indicate that this relationship is locus dependent. For position-sensitive loci, replication was most advanced when loci occupied their preferred nuclear environment and was altered when they occupied alternative locales, consistent with a functional coupling between nuclear position and replication timing. For these loci, efficient replication appears to depend in part on features provided or facilitated by the nuclear environment. In contrast, position-insensitive loci maintained similar replication timing across distinct nuclear environments despite having equally strong, or even stronger, canonical spatial preferences. This suggests that their replication program is maintained by mechanisms that are comparatively independent of current nuclear position. Thus, the coordinate establishment of nuclear position and replication timing at the TDP does not imply that position determines timing at every locus, but instead may reflect distinct position-dependent and position-independent modes of replication control. The preferential association of position-insensitive initiation zones with developmentally switching replication domains further suggests that, for these regions, replication timing may be established or reset through regulatory mechanisms that can operate largely independently of their immediate nuclear environment.

Sixth, RepTile also enabled genome organization to be compared across inferred cell-cycle stages. Compared with G1 cells, S phase cells showed stronger A/B compartment segregation, more stable chromatin-state-dependent radial positioning and reduced cell-to-cell variability in chromosome folding, indicating a more reproducible and compartmentalized configuration during S phase. These G1-to-S differences need not reflect restructuring initiated by S phase entry. Nuclear organization is re-established during early G1 around the TDP, and because G1 is unusually short in mESCs, the G1 population may contain cells at different stages of this reorganization. The more reproducible organization observed in S phase could therefore in part reflect completion of processes initiated before S-phase entry. In addition, replication-dependent histone gene clusters on different chromosomes showed increased spatial proximity specifically in S phase, together with a general increase in interchromosomal proximities between G1 and S phase. This organization may facilitate the coordinated expression or processing of genes required for DNA synthesis and chromatin assembly, although functional perturbations will be required to establish its role. More generally, these findings illustrate how retrospective cell-cycle staging can expose transient structural interactions that would be obscured when asynchronous cells are analyzed together.

Overall, our findings reveal distinct spatially dependent and locus-intrinsic modes of replication control and challenge the view that subnuclear position exerts a uniform influence on replication timing across the genome. Nevertheless, the analysis remains observational. Perturbations that reposition defined loci or disrupt specific nuclear environments will be required to distinguish among underlying regulatory mechanisms.

Several limitations define important directions for future work. RepTile currently infers replication from fixed-cell measurements and therefore reconstructs S-phase progression across a population rather than tracking the same locus through time. Its spatial associations cannot by themselves establish whether nuclear positioning is a cause or consequence of replication timing. Moreover, inference accuracy remains limited by sparse detection, creating a trade-off between confidence and genomic coverage. At 25kb resolution, approximately 70% of detected locus copies can be assigned a replication state at 82% accuracy, whereas increasing the accuracy threshold to 85% reduces coverage to approximately 55%. Further application of RepTile across diverse datasets, imaging platforms and species will be needed to assess its generalizability and enable comparative analyses of replication control. Finally, the balance between positional and intrinsic control may differ across cell types, developmental states and perturbations. Applying RepTile during differentiation, together with locus-repositioning experiments, nuclear-body perturbations and targeted deletion of candidate replication-control elements, should help establish the causal mechanisms underlying these two modes of regulation.

In summary, RepTile converts replication-dependent variation in multiplexed imaging data from a source of ambiguity into a directly measurable biological signal. By embedding replication states within the same three-dimensional nuclear context in which chromosome folding, nuclear landmarks and gene activity are measured, it enables a new class of single-cell structure–function analyses. The resulting observations challenge the idea that nuclear position broadly specifies replication timing and instead reveal locus-specific spatial and intrinsic modes of replication control, particularly at constitutive and developmentally regulated initiation zones.

---

**Extended Figure 1.**
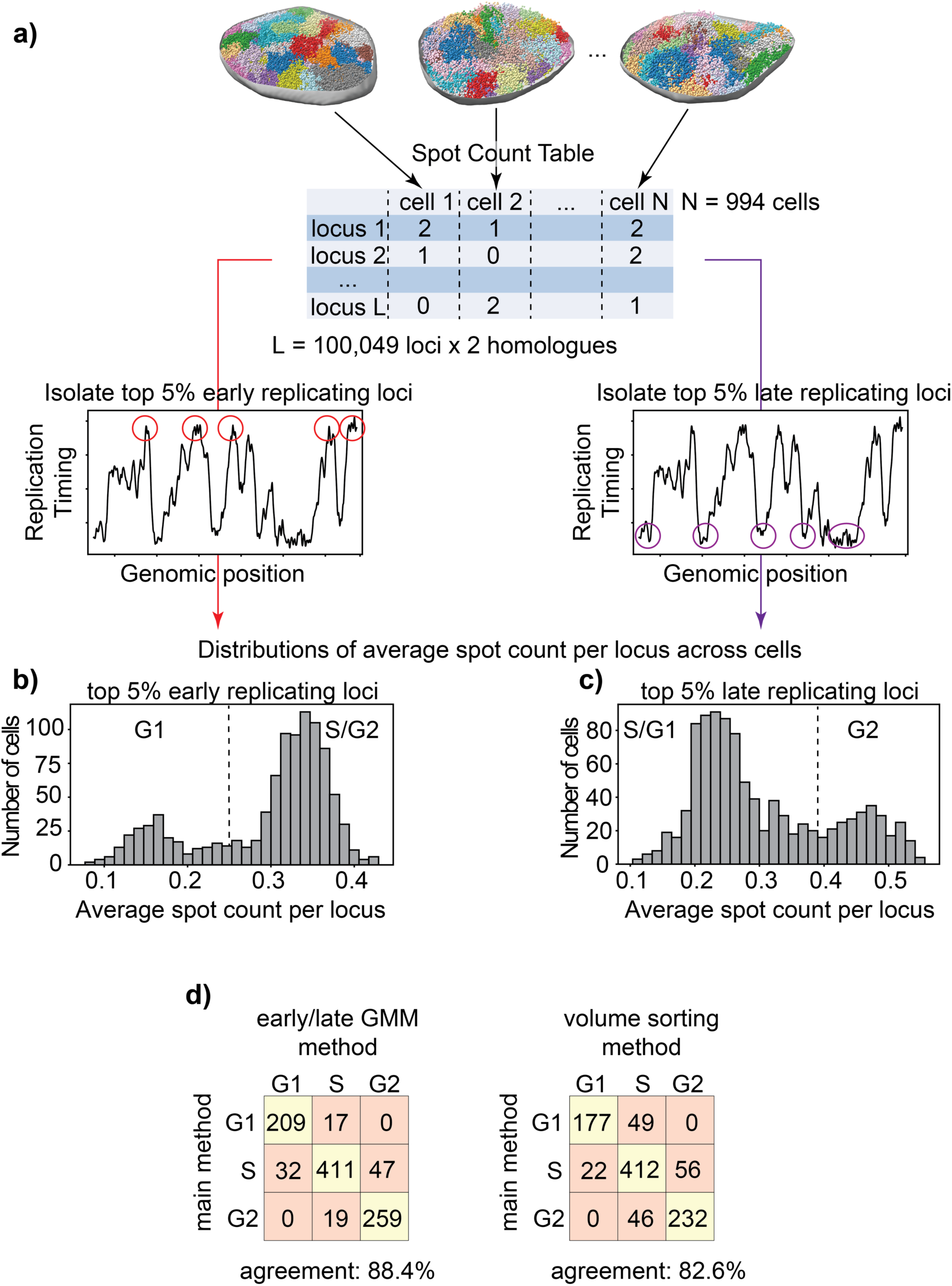
Alternative inference of single-cell G1, S and G2 labels. **a)** Workflow of the early/late Gaussian mixture model method (GMM) used to assign cells to G1, S and G2 phase (**Methods)**. Top: the spot-count distribution is determined for each cell. Bottom: The earliest-replicating 5% (left) and latest-replicating 5% (right) loci are identified from the replication timing (RT) signal ^38^ and average spot count for each locus set is calculated for each cell. **b-c)** Distributions, across cells, of the single-cell average spot count for the earliest-replicating (**b**) and latest-replicating (**c**) loci. Dashed lines indicate the GMM classification thresholds, and the inferred cell-cycle stages are indicated (**Methods**). **d)** Confusion matrices comparing cell-cycle assignments from the primary method with those from the two alternative methods, namely early/late GMM and nuclear volume sorting (**Methods**). Overall agreement is reported below each matrix.

**Extended Figure 2.**
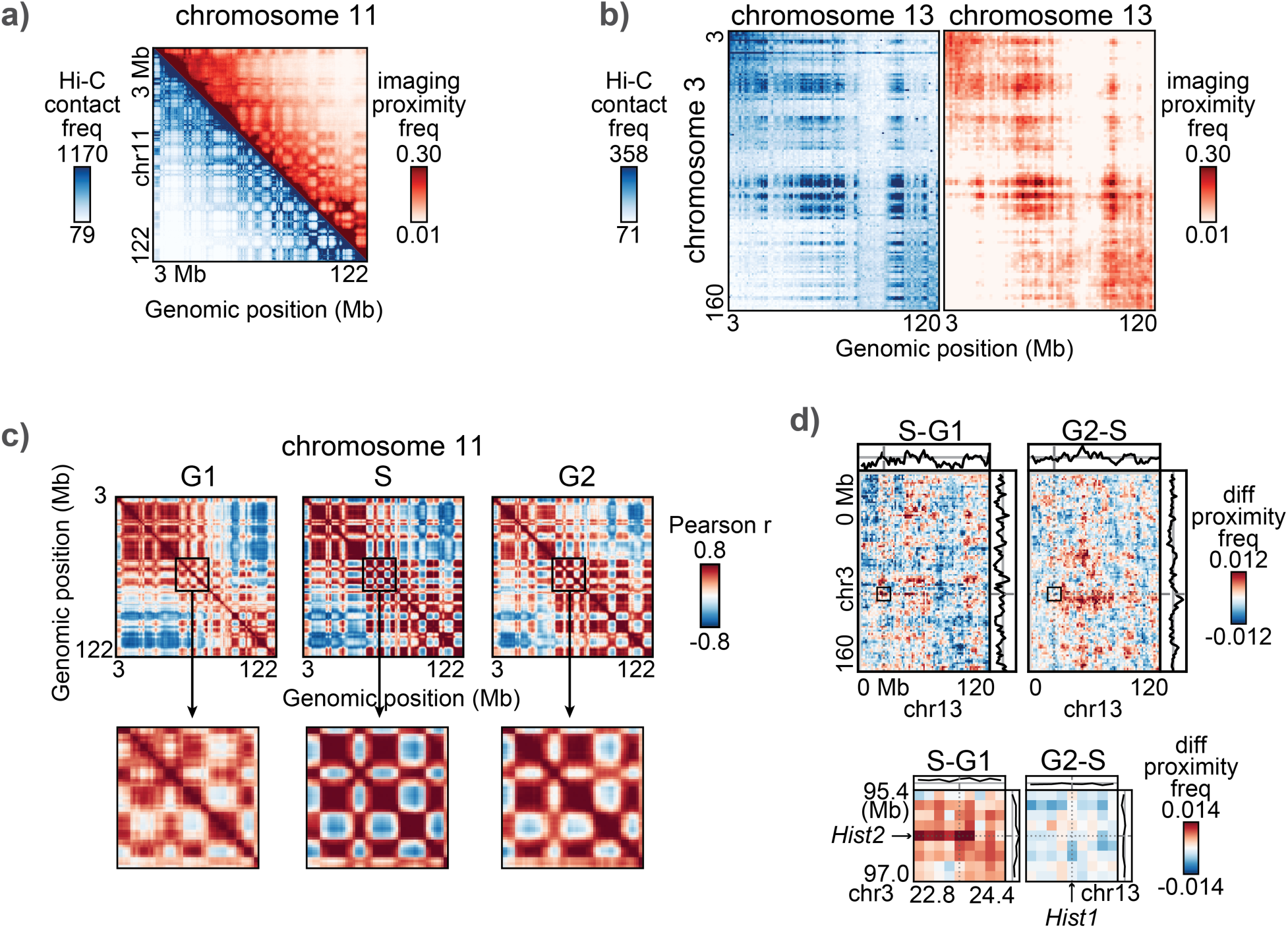
Imaging-derived proximity frequency matrices agree with Hi-C contact frequency maps and resolve cell-cycle-specific chromatin interactions. a-b) Hi-C contact frequency maps (shown in blue) ^34^ and imaging-derived proximity frequency matrices (shown in red), averaged across all cells for intra-chromosomal maps for chromosome 11 at 250kb resolution (**a**, Pearson r = 0.91) and inter-chromosomal maps between chromosomes 3 and 13 at 1Mb resolution (**b**, Pearson r = 0.73). **c)** Row-wise correlation maps for G1, S and G2 from the imaging-derived proximity frequencies (**Methods**). A 25Mb region is shown in higher magnification. **d)** Inter-chromosomal S-G1 and G2-S differential proximity frequency maps for chromosome 3 vs chromosome 13 at 1.5Mb resolution (top) and a zoomed 1.8Mb region spanning the *Hist1*-*Hist2* gene cluster interaction at 200kb resolution (bottom).

**Extended Figure 3.**
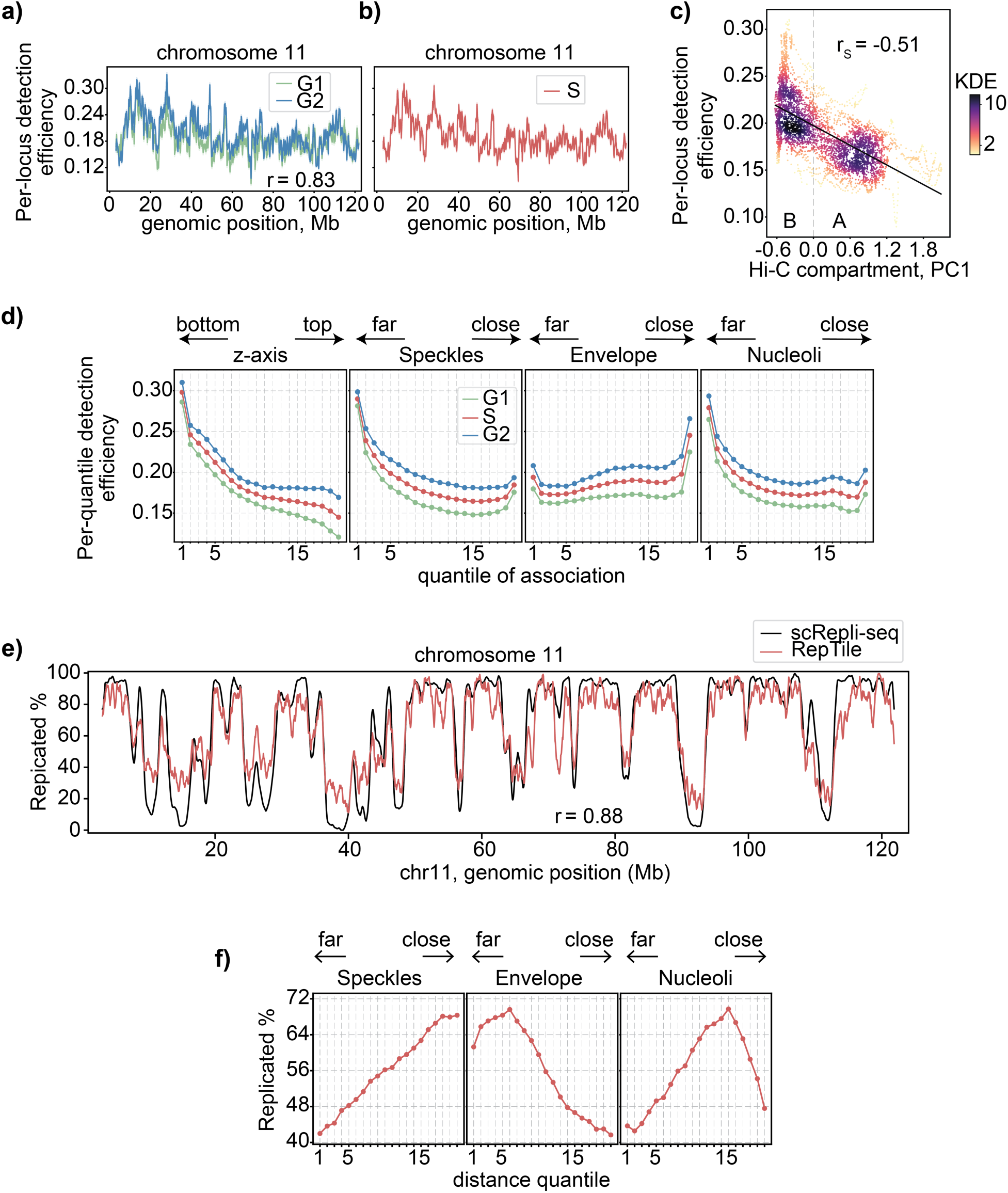
Detection biases and validations of RepTile’s statistical module. a-b) Mean locus-dependent detection efficiency for chromosome 11, inferred separately from G1-and G2-phase cells (**a**) and for S cells (**b**, **Methods**). The Pearson correlation between G1-and G2-derived signals is indicated. **c)** Scatter plot of locus-specific mean detection efficiency in S-phase cells against Hi-C A/B compartment PC1 score ^37^. The dashed line represents the linear regression fit. The vertical dotted line indicates the separation between A compartment (PC1 > 0) and B compartment (PC1 < 0). The Spearman correlation is indicated. **d)** Average detection efficiencies across quantiles defined by nuclear envelope distance, local speckle-and nucleolar-marker intensities, and z-axis position, shown separately for cells in G1-, S-and G2-phase (**Methods**). **e)** Locus-specific average replicated fractions across chromosome 11 measured by single-cell Repli-seq ^39^ at 50kb resolution (black) and inferred by RepTile at 25kb resolution (red, **Methods**). The Pearson correlation is indicated. **f)** Percentage of replicated loci across quantiles defined by nuclear-envelope distance and local speckle-and nucleolar-marker intensity, averaged across S-phase cells.

**Extended Figure 4.**
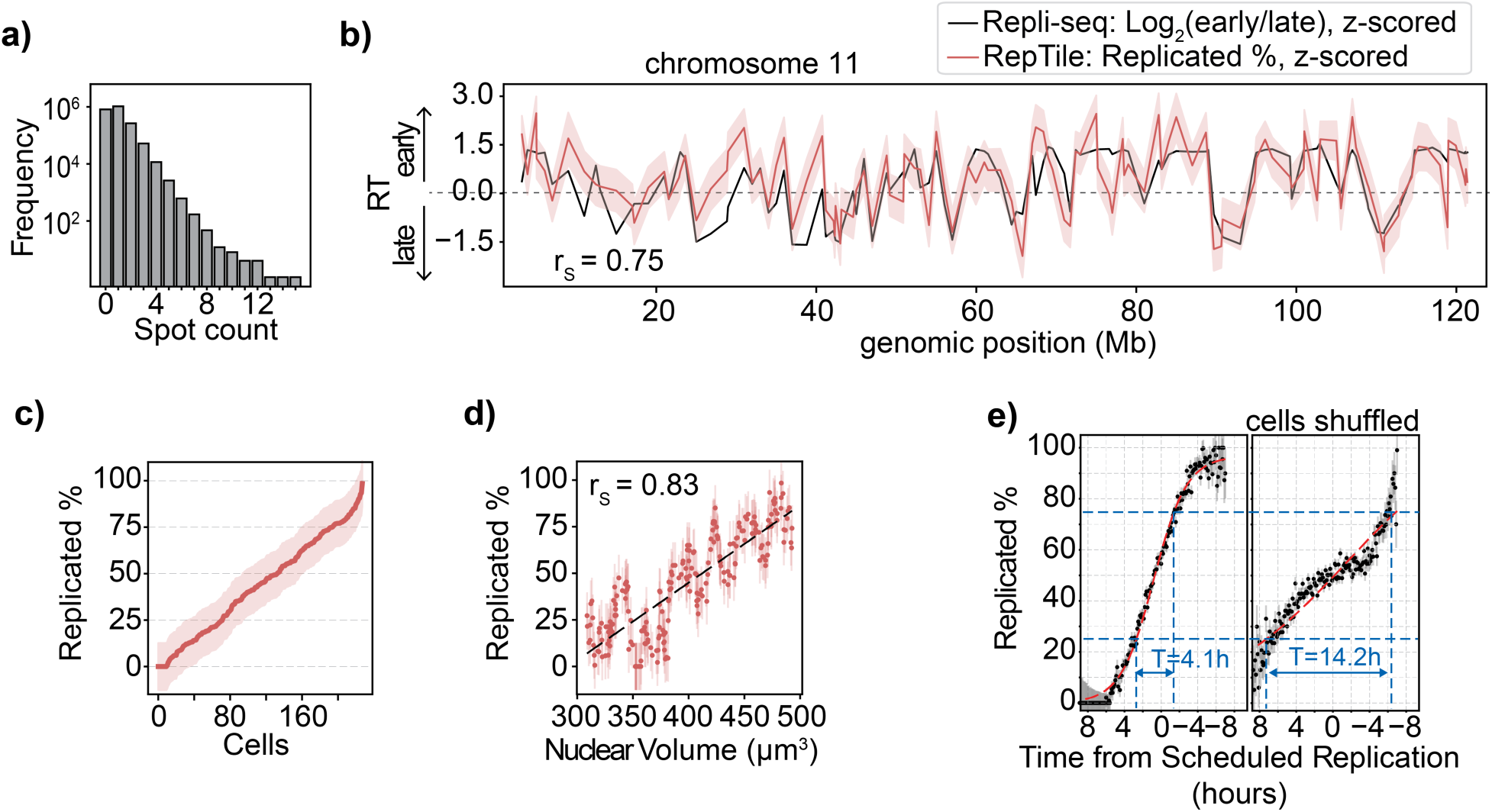
Validation of RepTile using low-coverage DNAseqFISH+ data. **a)** Distribution of spot counts per locus allele across all cells in the low-coverage DNAseqFISH+ dataset, comprising 2,460 loci sampled at intervals of approximately 1Mb^19^. **b)** Replication timing (RT) inferred by RepTile (red) and by Repli-seq (black) ^38^. Dashed shading indicates error uncertainty in RepTile estimates and the Spearman correlation is indicated. **c)** Percentage of replicated DNA inferred by RepTile in individual S-phase cells. Cells are ordered by their percentage of inferred replicated DNA and dashed shading indicates uncertainty. **d)** Relationship between inferred percentage of replicated DNA and nuclear volume in each S-phase cell with vertical bars indicating error bars in inferred replicated DNA, and a dashed black line indicating the linear fit. **e)** Percentage of inferred replicated DNA (with error bars) calculated for each 0.1-hour bin of the Time from Scheduled Replication (left, **Methods**), with the analogous curve obtained after randomly shuffling the S-phase cell ordering (right). The red curves indicate sigmoid fits, while the blue dashed lines show the time required for the replicated fraction to increase from 25% to 75%. Blue lines indicate the resulting T-width.

**Extended Figure 5.**
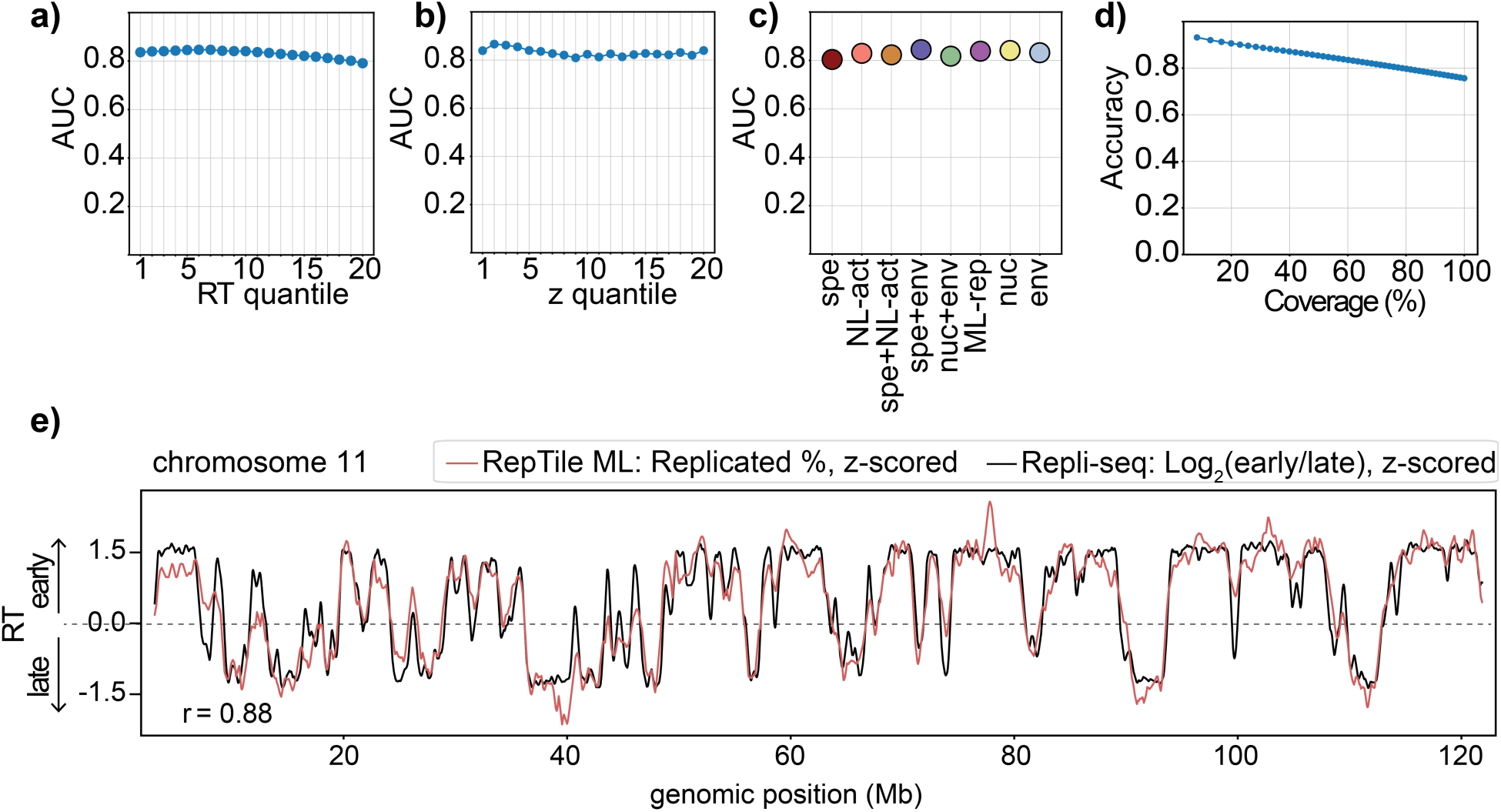
Performance and validation of RepTile’s machine-learning module. a-b-c) Area under the receiver operating characteristic curve (AUC) for RepTile’s machine-learning (ML) module (**Methods**), averaged across chromosomes and stratified by replication-timing (RT) quantiles (**a**), z-axis quantiles (**b**) and nuclear locales (**c, Methods**). **d)** RepTile ML’s classifier accuracy (y-axis) as a function of coverage (x-axis), defined as the fraction of loci with assigned replication state. Each point uses a different classification threshold to binarize the classifier output (**Methods**). **e)** Replication-timing signal along chromosome 11 measured by Repli-seq ^38^ (black) and derived by averaging RepTile ML’s predicted replication states across cells (red, **Methods**). Both signals were z-scored, and their Pearson correlation is indicated.

**Extended Figure 6.**
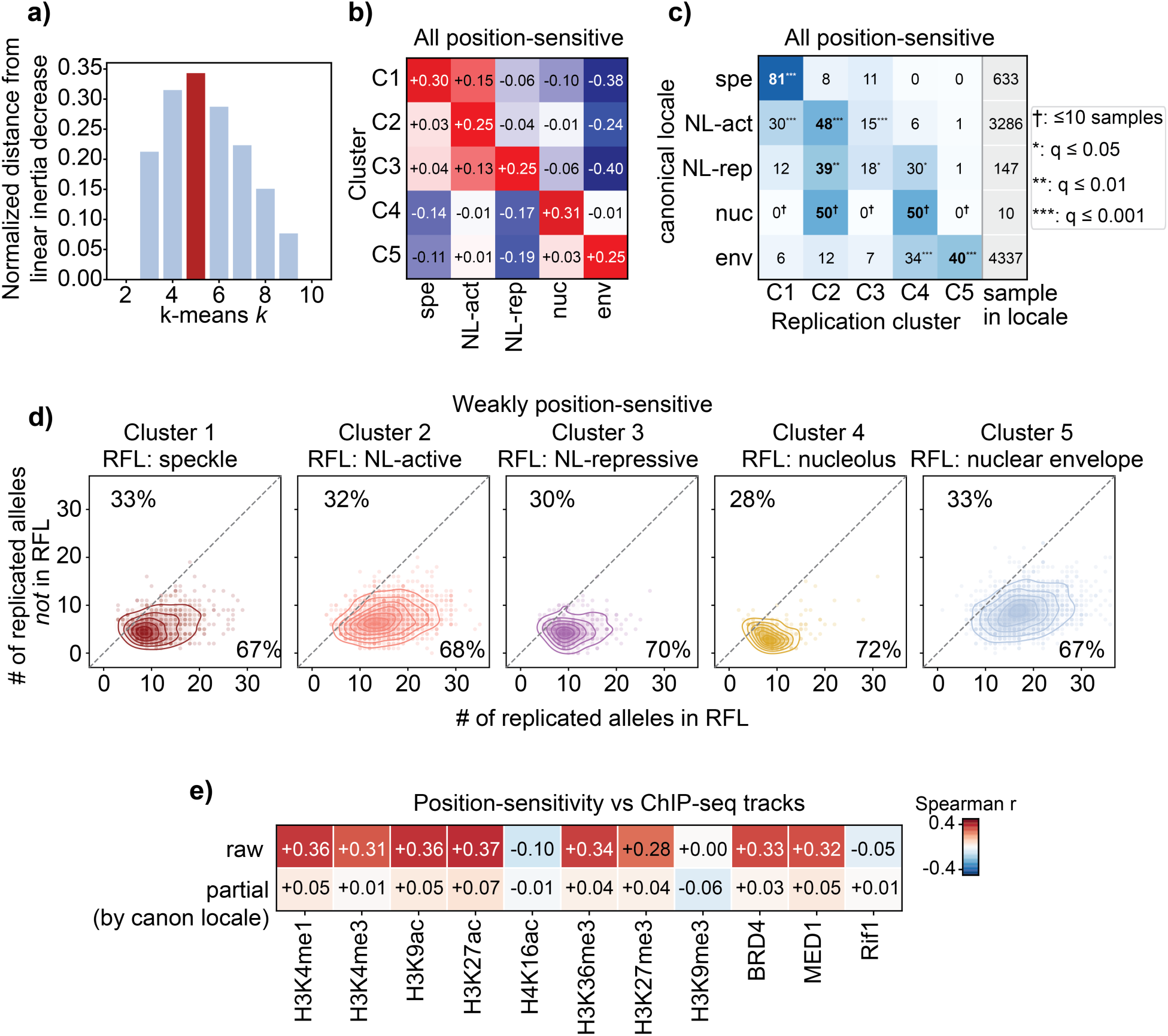
Clustering and characterization of position-sensitive loci. **a)** Selection of the optimal number of replication-locale clusters (*k*) using the Kneedle method ^58^ for position-sensitive loci using k-means clustering. For each *k* (x-axis), the method measures the perpendicular distance between the min-max normalized inertia (within-cluster sum of squares) and the linear interpolation between the minimum and maximum (y-axis, **Methods**). The maximum distance identified *k* = 5 as the optimal number of clusters (red). **b)** Mean locale-associated replication shift Δ*RF* (locale), across loci in each cluster and for each primary locale, considering all position-sensitive loci (**Methods**). For each locus *i*, Δ*RF_i_* (locale) is defined as the difference between the fraction of replicated instances when the locus occupied a given locale and the fraction replicated when it occupied any other locale. **c)** Confusion matrix showing the percentage of loci with each canonical locale (rows) assigned to each of the five replication-locale clusters (columns) for all position-sensitive loci. Rows are normalized to 100%, and the sixth column indicates the total number of loci for each canonical locale. Bold values indicate the largest percentage in each row. Stars indicate Benjamini-Hochberg-corrected q-values from one-sided permutation tests for over-representation relative to a null model of independence between canonical locale and cluster (*q < 0.05, **q < 0.01, ***q < 0.001, **Methods**). Daggers (†) indicate percentages calculated from fewer than ten loci with the nucleolus canonical locale. **d)** Per-locus scatter and contour plots for all weakly position-sensitive loci, showing the number of discordant-allele cells, in which the replicated allele occupied the locus’s replication-favorable locale (RFL, x-axis, **Methods**) or another locale (y-axis). The dashed line indicates equal frequencies. Percentages below and above the y = x line indicate the fractions of allele-discordant cells in which the replicated allele occupied the RFL or another locale, respectively. Loci are grouped by replication-locale cluster (**Methods**). **e)** Spearman correlations (top) and partial Spearman correlations controlling for canonical locales (bottom) between position sensitivity *s* and ChIP-seq tracks of several epigenetic markers (200kb, **Methods**).

**Extended Figure 7.**
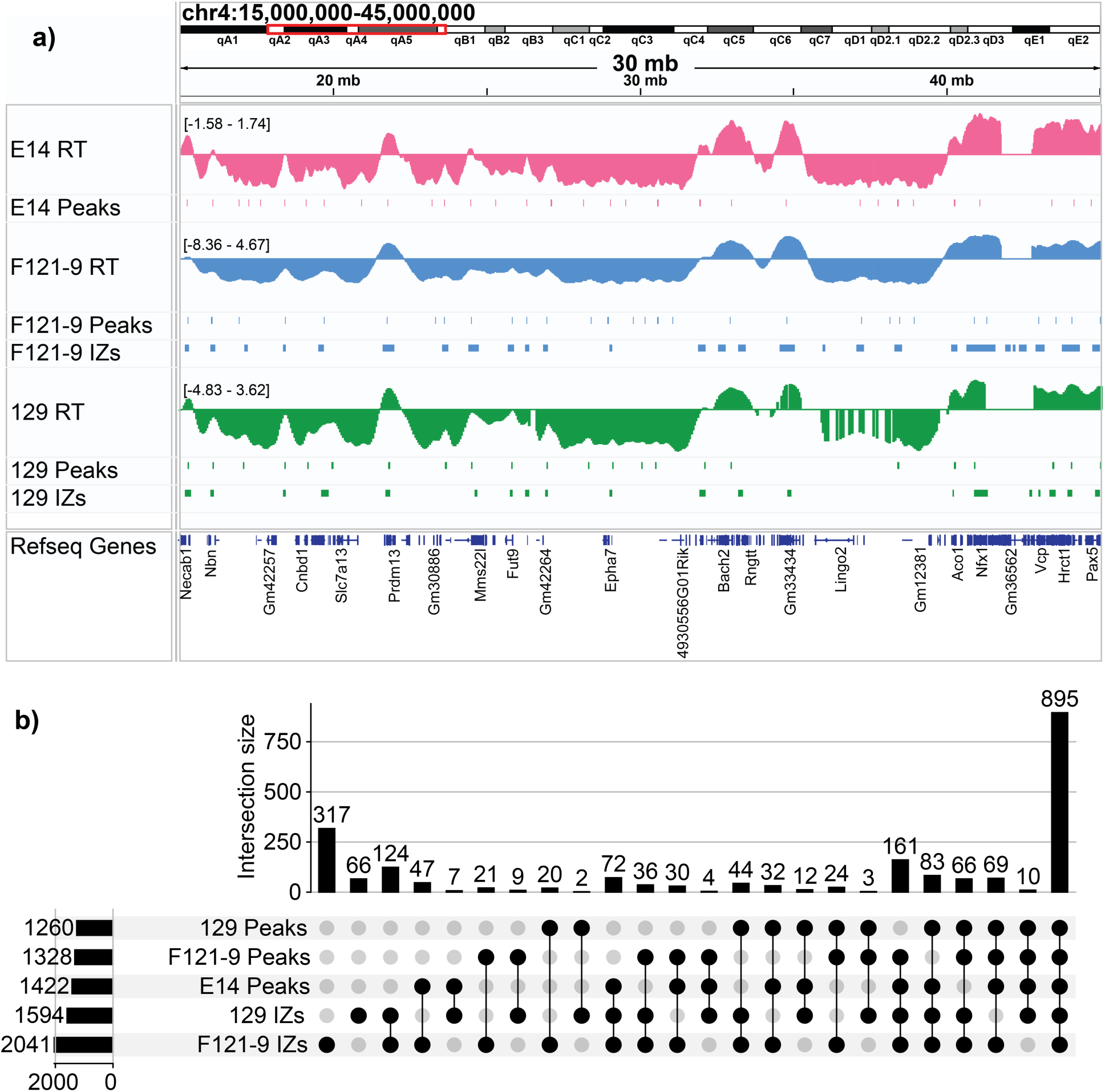
Selection of E14 initiation zones from intersected Repli-seq datasets. **a)** Plots of eight mESC replication signals across genomic positions for a 30Mb area on chromosome 4 (chr4:15-45Mb). From top to bottom the tracks are: 1) E14 RT from Repli-seq ^38^, 2) peaks identified from the E14 RT (**Methods**), 3) F121-9 RT ^59^ and 4) peaks identified from F121-9 RT, 5) F121-9 initiation zones (IZs) from high-resolution Repli-seq ^40^, 6) parsed 129 RT from F121-9 ^59^ and 7) peaks identified from it, 8) parsed 129 IZs from F121-9 ^40^. This plot also indicates some gene positions. **b)** UpSet plot ^60^ (**Methods**) obtained from the following regions: 1) E14 RT peaks, 2) F121-9 RT peaks, 3) F121-9 IZs, 4) 129 RT peaks, 5) 129 IZs. The largest intersection (N=895) is shared by five sets, and defines the selected set of E14 initiation zones.

## Methods

### Imaging datasets

We applied our approach to two different datasets. The first is a *DNAseqFISH+* dataset of *E14 mouse embryonic stem cells (mESC)* ^16^. The experiment targeted 100,049 non-overlapping 25kb genomic regions tiled across the mappable mouse genome. Unmappable regions, including centromeres and telomeres, were not imaged. Across the 1,076 imaged cells, the dataset provides three-dimensional (3D) coordinates for over 64 million detected fluorescent spots, each uniquely assigned to one of the targeted genomic regions. The dataset also reports, for each spot, the fluorescence intensity values of 65 sequential immunofluorescence antibodies, measured in the voxel occupied by the spot and z-scored across voxels per marker and per cell. For markers enriched in sub-nuclear bodies, such as SF3A66 at nuclear speckles and Fibrillarin at nucleoli, these intensities provide proxies for the spatial proximity of each spot to the corresponding body. We refer to this data set as E14-2025.

The second is an independent *DNAseqFISH+* dataset from *E14 mESC* ^19^, with 2,460 genomic regions imaged across 446 cells. This dataset sampled the genome more sparsely, targeting 25kb regions at intervals of approximately 1Mb. We refer to this data set as E14-2021.

### Quality control for single cells

For the E14-2025 dataset, out of the 1,076 cells, 71 were discarded due to being extreme outliers in nuclear volume or total spot counts (below the 2^nd^ or above the 98^th^ percentiles), or being extreme outliers with respect to the observed spot counts and expected spot counts predicted by the linear regression of spot count on nuclear volume (outside the 0.5-99.5 percentile range of that ratio). 11 cells were removed because they displayed clear distortions on visual inspection, leaving a total of 994 cells.

### Separation of homologous copies

The two homologous copies of each autosomal chromosome were separated as previously described for DNAseqFISH+ ^19^. In each cell and for each autosome, loci were first separated into two clusters by Ward clustering ^61^. If the two clusters were not well separated, they were re-partitioned by spectral clustering ^62^ on the affinity matrix *A*_*ij*_ = *exp*(−*D*_*ij*_/*δ*), where *D_ij_* is the Euclidean distance matrix and *δ* its standard deviation. Clusters were considered well separated when the distance between their centroids exceeded 1.2 times the sum of their spreads, defined as the standard deviation of the loci within each cluster. When two or more spots of the same locus were assigned to the same copy, they were removed if more than 5μm apart.

### Coarse-graining structures and linear interpolation

For the E14-2025 data we generated structures at 25kb, 200kb, 500kb and 1.5Mb for visualization and analysis. We tiled the genome in regularly spaced bins at the target resolution and assigned each bin the center of mass of all overlapping 25 kb loci. Immunofluorescence marker values were coarse-grained by averaging over the same loci.

To infer the nuclear locations of undetected regions we linearly interpolated position and marker values to the nearest imaged loci to their left and right. When no left or right locus was available, the missing locus was assigned the position of the closest imaged locus along the genomic chain.

### Estimation of nuclear envelope

The nuclear envelope of each cell was estimated by fitting an α-shape ^63^ to all imaged loci, using α = 0.5μm^-1^ for the E14-2025 data and α = 0 for the E14-2021 data, which reduces the α-shape to a convex hull ^64^. The convex hull was also used for E14-2025 visualizations. Distances between a 3D point and the envelope were computed as the Euclidean distance to the closest point on the envelope.

### Definition of nuclear locales

For the E14-2025 dataset, we assigned each 25kb locus in each cell to one of eight nuclear locales from its subnuclear position and imaged chromatin marks. Speckle, nucleolar and envelope associations were defined using SF3A66, Fibrillarin and envelope proximity, respectively. Within each cell, loci were called landmark-associated if their signal was above a per-landmark percentile cutoff. These cutoffs were selected by optimizing the correlation between population-average association profiles and bulk SON, MKI67 or LaminB1 TSA-seq ^27^. The resulting cutoffs were the 95^th^ percentile for nucleoli (top 5%, r = 0.75), the 60^th^ for the envelope (top 40%, r = 0.87) and the 85^th^ percentile for speckle (top 15%, r = 0.84). Loci passing two cutoffs were assigned to the corresponding mixed locale. Those passing none were classified by whether their strongest mark was active (RNAPII-Ser2P/Ser5P, H3K4me1/2/3, H3K9ac, H3K14ac, H3K27ac, H4K8ac or H4K16ac) or repressive (H3K9me2/3, H3K27me3 or H4K20me3). Each locus was thereby assigned uniquely to speckle, nucleolus, envelope, one of three mixed locales, non-landmark active or non-landmark repressive.

### Canonical locale, canonical strength and locale entropy

We defined the ‘canonical locale’ as the per-locus most frequent locale across all cells and allele-copies, and the ‘canonical strength’ as the ratio of its frequency to that of the runner-up. Locale entropy was the normalized Shannon entropy:

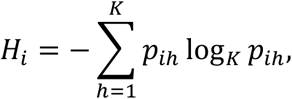

where *p_ih_* is the frequency of locus *i* in locale *h* and *K = 8*. Assuming multinomial covariances, uncertainty was estimated by the delta method:

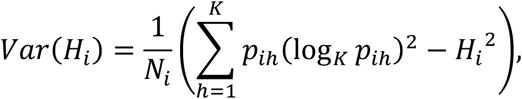

where *N_i_* is the number of cell-copy instances in which locus *i* was detected.

### Identification of nuclear bodies by empty nuclear volumes

For 3D visualization, we imputed speckle, nucleolar and pericentromeric and telomeric heterochromatin volumes as regions depleted of DNA. For each cell, we voxelized the nucleus at 200nm resolution and computed a Gaussian kernel density estimate (KDE) over all imaged loci. Subtracting this map from its maximum produced a negative-density map. Voxels within 750nm of the envelope were excluded to avoid detecting the locus-sparse nuclear periphery. Low-density regions were classified using SF3A66 for speckles, Fibrillarin, rDNA, Rnu3b_RNA and ITS1_RNA for nucleoli, and MajSat, MinSat and Telomere for heterochromatin. In each cell and for each body, loci above the 80^th^ percentile of the maximum signal across its markers defined a body-specific KDE. Each voxel was assigned to the body with the highest KDE, separating the negative-density map into body-specific maps. Finally, we retained voxels above the 99.5^th^ (nucleoli) or 99^th^ (speckle) percentile of the body-specific map, and applied one round of erosion followed by two (nucleoli) or one (speckles) rounds of dilation. Only speckles and nucleoli were rendered in 3D.

### Infer cell-cycle stage for each cell

For the E14-2025 dataset, we inferred cell-cycle labels by maximum-likelihood fitting of per-cell total spot counts, *X*. We modelled *X = M + E*, where *E* is a zero-mean Gaussian variability term, *M* is a copy-number contribution (2C in G1, 4C in G2, and increases uniformly in S). The per-cell total spot count probability densities were:

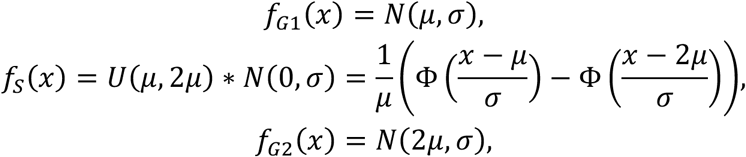

where *N(μ, σ)* is the Gaussian distribution with mean *μ* and standard deviation *σ*, *U(μ, 2μ)* is the uniform distribution between *μ* and *2μ*, ∗ denotes convolution, and *Φ* is the standard normal cumulative distribution function. Indicating the unknown phase fractions as *π_k_*, the per-cell density and independent-cell log-likelihood were:

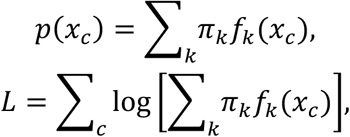

with the constraints:

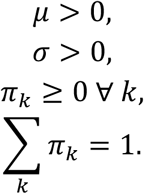

We minimized *−L*, enforcing the constraints via reparameterization, obtaining the best estimates *µ̂*, *σ̂*, *π̂*_*G*1_, *π̂*_*S*_, *π̂*_*G*2_, with the *π* values providing the population fractions of G1, S and G2.

To assign a cell with total spot count *x_c_* to a cell-cycle stage, we calculated the posterior phase probabilities:

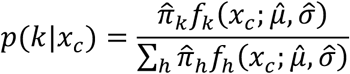

and assigned the top *π̂*_*S*_ fraction cells ranked by *p(S|x_c_)* to S phase and the remaining cells to G1 or G2 by the larger posterior.

### Additional methods to infer cell-cycle labels

We assessed robustness using two additional methods (**Extended Figure 1abcd**).

#### Gaussian mixture on early/late loci

We defined early and late loci as the top and bottom 5% of Repli-seq RT ^38^ and, per cell, averaged their spot counts. Two-component Gaussian mixture models were fitted separately to both distributions. The lower-count early-locus component defined G1 and the higher-count late-locus component defined G2. Remaining cells were assigned to S phase.

#### Volume sorting and FACS percentages

We ranked cells by nuclear volume and used external FACS fractions to estimate the percentage of G1, S, G2 ^33^. The lowest-volume x_G1_% and highest-volume x_G2_% were assigned to G1 and G2, respectively, and the remainder to S phase. This method was also used for the E14-2021 dataset.

### Construction of intra-and inter-chromosomal proximity frequency maps

We generated proximity frequency maps from how often a pair of loci fell within 1μm. Each chromosomal copy was treated independently: the number of valid samples was 2n_cells_ for intra-chromosomal maps (ignoring inter-copy interactions) and 4n_cells_ for inter-chromosomal maps. For each sample *c*, we calculated a proximity tensor *L_cij_*, equal to 1 when bins *i* and *j* were within 1μm in sample *c* and 0 otherwise. We also calculated a co-presence tensor *N_cij_* to prevent missing-locus bias.

After averaging L and N across samples to obtain *l_ij_* and *n_ij_*, proximity frequency and its delta-method variance were:

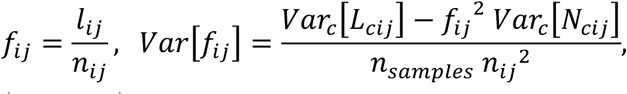

Where we used *cov*_*c*_(*L*_*cij*_, *N*_*cij*_) = *f*_*ij*_*Var*_*c*_[*N*_*cij*_], obtained by modeling contacts as binomial in co-presence events. Maps were calculated at 200kb and 1.5Mb bin windows. For the latter, proximities were first detected at 100kb and then aggregated. Maps were generated for G1, S, G2 or all cells. Because of nuclear expansion throughout the cell cycle, small G1 nuclei had inflated proximity frequencies compared to S and G2. Thus, we rescaled each state’s mean intra-and inter-chromosomal frequency map by one scalar to match the corresponding all-cell mean.

To determine intra-chromosomal proximity frequency per sequence distance for each state, we pooled 200kb intra pairs genome-wide by genomic separation and calculated their mean contact frequency *Φ(|i−j|)*, from 200kb to 120Mb. Curves were smoothed with a five-bin (±400kb) window.

### Comparison of imaging-derived proximity frequencies with contact frequencies from Hi-C data

We compared imaging maps with Hi-C ^34^: we extracted Knight-Ruiz-normalized matrices at the imaged bins and clipped at the 6^th^-94^th^ percentiles for intra maps and 1^st^-99^th^ for inter maps. Pearson correlations were calculated for each intra-(250kb) and inter-chromosomal (1Mb) map and averaged. For intra maps, we also calculated genomic-distance-adjusted Pearson correlations by averaging correlations along each off-diagonal, discarding distances with fewer than 100 valid pairs.

### Compartment analysis from contact frequency matrices

We called compartments at 200kb, per-chromosome and per-state ^65^. Proximity maps were normalized by genomic distance averages, yielding the observed / expected maps: *OE_ij_ = f_ij_ / Φ(|i−j|)*. OE rows were z-scored, diagonal and missing entries set to zero, and used to calculate the pairwise correlation matrix *R_ij_*, which was then symmetrized. We applied Principal Component Analysis to each locus, retained among the first five components those explaining at least 5% of variance, and selected the component most correlated in absolute value with gene density as the compartment component. This eigenvector was L2-normalized and oriented to correlate positively with gene density.

For saddle plots, loci were ranked by eigenvector value, divided into 50 equally occupied quantiles per chromosome, and the OE was averaged between quantile pairs across chromosomes, excluding contacts below 1 Mb. Compartment strength was calculated as *[mean(OE_AA_) + mean(OE_BB_)] / mean(OE_AB_)*, defining A and B as the top and bottom 5% of values within each chromosome.

### Expected inter-chromosomal interactions per locus

For each 200kb locus, we calculated its expected number of inter-chromosomal proximities by summing its inter-chromosomal frequencies, *τ*_*i*_ = ∑_*j*: *chr*(*j*)≠*chr*(*i*)_ *f*_*ij*_. State-specific gain was Δ*τ*_*i*_(*s*) = *τ*_*i*_(*s*) − max(*τ*_*i*_(*s*^’^), *τ*_*i*_(*s*^’’^)), and was normalized by max(*τ*_*i*_(*s*^’^), *τ*_*i*_(*s*^’’^)) to convert it into a percentage.

### Analysis of trans to total ratio

For each locus in each cell, we computed the trans-to-total frequency *ρ* as the fraction of loci within radius *r* belonging to another chromosome or homolog. To visualize distributions, *ρ* was averaged per-cell across genome-wide loci at 25kb with *r* = 500nm, together with the coefficient of variation (CV) of *ρ* across cells (standard deviation divided by mean) computed per-locus separately in G1, S and G2 cells. The significance of the changes in median CV across states was assessed with a permutation test, shuffling cell-cycle labels 10,000 times. To identify S-specific loci, *ρ* was averaged per-locus across G1, S and G2 cells at 200kb with *r* = 1μm, and scored as Δ*ρ*_*i*_ = *ρ*_*i*_(*S*) − max(*ρ*_*i*_(*G*1), *ρ*_*i*_(*G*2)). Significance was again assessed by permutation, randomly shuffling cell-cycle labels 10,000 times, correcting the two-sided p-values with Benjamini-Hochberg ^66^, calling significant loci at FDR q ≤ 0.05.

### Hist1/Hist2

We annotated the location of the *Hist1* (chr13:21.6-22.2Mb + chr13:23.4-24.0Mb) and *Hist2* (chr3:96.2-96.4Mb) replication-dependent histone gene clusters, identified the 200kb regions mapping to them, and calculated the average proximity frequency among all pairs of loci mapping their interaction, separately in G1, S and G2. We compared states with two-sided z-tests of the S-G1 and G2-S differences, correcting the standard error for covariance between bin pairs.

### Envelope distance profile analysis

We calculated per-locus nuclear envelope distance profiles for G1, S and G2. To correct for nuclear expansion, distances were quantile-normalized within each cell from 0 (furthest) to 1 (closest) and averaged by state, thus measuring radial genome organization. For S-G1 and G2-S, differences were tested using two-sided Mann– Whitney U tests on per-cell normalized distances, and p-values were corrected within each transition by Benjamini-Hochberg, with significance at FDR q ≤ 0.05. We also calculated each locus’s coefficient of variation by state.

### RepTile framework

RepTile is a bi-modular framework that infers replication directly from multiplexed FISH imaging and chromosome tracing experiments. Its statistical module uses a parametric model to infer the fraction of replicated DNA in any selected subset of loci, whereas its machine-learning module classifies individual loci in single cells as ‘replicated’ or ‘unreplicated’.

### RepTile’s statistical module

This module infers the fraction of replicated DNA in any selected subset of loci, within the same cell or across different cells. The input is the spot-count distribution across the selected locus-cell pairs, with homologous copies treated as separate locus instances. We modeled the observed spot count *N* as a function of replication state, detection efficiency and overcounting

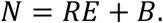

*R* is the replication state, defined as *R = 1* for an unreplicated locus and *R = 2* for a replicated locus, modeled as a Bernoulli variable with replication probability *p*. *E* models detection with efficiency *ε*. An unreplicated locus is detected with probability *ε*, while for a replicated locus, both, one or neither sister copies are detected with probabilities *ε²*, *2ε(1 – ε)* and *(1 − ε)²*, respectively. Thus, *P*(*E* = *e* | *R* = *r*) = *Binomial*(*re*; *r*, *ε*). B models spurious splitting of spots representing a single locus into multiplets, i.e. “overcounting” events: each detected sister can generate overcounting, with *P*(*B* = *b* | *R* = *r*, *E* = *e*) = *Poisson*(*b*; *reβ*).

We inferred *p*, *ε* and *β* using the Generalized Method of Moments (GMM) ^67^, by matching the observed average spot count and zero-count fraction *f* to the model predictions:

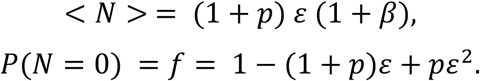

Because these two moments cannot determine all three unknowns, we used G1-and G2-phase cells, for which *p_G1_ = 0* and *p_G2_ = 1*, to calibrate *ε* and *β*. With these parameters fixed, we then inferred *p* in S-phase cells. G1/S/G2 labels can either be obtained experimentally or inferred as described above.

### Estimating S-phase progression in single cells

We sorted S-phase cells according to their per-cell fraction of replicated DNA. Detection parameters were first estimated independently in each G1-and G2-phase cell using all genome-wide loci, excluding chromosome X because of its different ploidy. For each G1/G2 cell *c*, we measured the average spot count, <*N*>_*c*_, and the fraction of zero counts, *f_c_*, and estimated the cell-specific *ε_c_* and *β_c_* from the RepTile model:

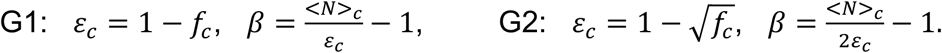

Remarkably, the average detection efficiency values across G1-and G2-phase cells were similar (0.18 and 0.20, respectively), with a relatively low coefficient of variation (7.2%). We therefore approximated the detection efficiency of each S-phase cell by the average of G1-and G2-phase estimates, 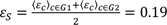, and propagated the corresponding standard deviations as an estimation of error uncertainty. To avoid bias from cells with artificially low detection efficiencies at the smallest nuclear volumes, only G1/G2 cells with nuclear volumes above 400μm³ for E14-2025 and 210μm³ for E14-2021 were included in this calibration. We then inferred for each S-phase cell the replicated fraction *p_c_* and overcounting parameter *β_c_* as

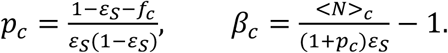

Because GMM estimates are not intrinsically constrained to the parameter bounds, *p_c_* was clipped to the interval [0, 1]. Error uncertainties in *p_c_* and *β_c_* were propagated through the GMM equations using the delta method.

For the lower-coverage E14-2021 dataset, we increased the statistical power of the single-cell inference by first ordering cells according to their nuclear volume as an initial first proxy estimate of cell-cycle progression. We then calculated the average spot count per cell, <*N*>*_c_*, and the fraction of zero counts, *f_c_*, using a sliding window smoothing, pooling the data of n=11 consecutive cells (central ± 5 cells), and then applied RepTile as described above.

### Estimating locus-specific replication timing

For each locus *i*, its replication timing (RT) was estimated as the fraction of S-phase allele instances in which the locus was replicated. To calculate this quantity, we first estimated locus-specific detection efficiency *ε_i_* and overcounting parameter *β_i_* from G1-and G2-phase cells. For each locus, we calculated the average spot count, <*N*>*_i_*, and the fraction of zero counts, *f_i_*, separately in G1 and G2 cells, and then used RepTile’s equations:

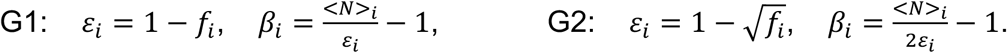

Because G1 and G2 locus-dependent efficiencies were highly correlated (Pearson r = 0.83, p-value < 1e-300), the S-phase efficiency for each locus was estimated as their signal-average, *ε*_*i*_^*S*^ = (*ε*_*i*_^*G*^^1^ + *ε*_*i*_^*G*^^2^)⁄2. We then inferred *p_i_* and *β_i_* across S-phase cells, clipped *p_i_* to [0, 1], and propagated errors through the GMM equations using the same formulas as in the previous paragraph.

As a negative control, we estimated RT without RepTile by simply averaging the spot count across S-phase allele instances for each locus. Genomic correlations at 25kb were calculated with a sliding window smoothing of 300kb.

### Reproducing replication timing heterogeneity (T-width)

To quantify cell-to-cell replication-timing heterogeneity, we adapted the T-width framework introduced for single-cell Repli-seq ^39^. We placed loci and cells on a common 10-hour S-phase axis. Locus RT values were inverted, clipped to the 0.01^th^ and 99.99^th^ percentiles, then rescaled to [0h, 10h] to obtain the scheduled replication time of each locus, *T_i_*. Then, for each S-phase cell (*c*) the inferred replicated fraction was multiplied by 10h to obtain its elapsed S-phase time *T_c_*. We finally calculated the ‘time from scheduled replication’ for locus *i* in cell *c* as *TSR_ic_ = T_i_ − T_c_*.

We binned TSR from −10h to 10h in 0.1h intervals and estimated the replicated fraction in each bin. Each *TSR*-bin therefore contains a unique composition of loci in S-phase cells such that the difference *T_i_ − T_c_* is constant within 0.1h. The estimation of detection efficiency required G1 and G2 data with the same locus composition. However, since *TSR* is not defined for G1 and G2 cells, we implemented a bootstrap approach. We randomly paired each S-phase cell contributing to the *TSR*-bin with one G1 and one G2 cell, transferring its cell-specific composition of loci. Thus, all TSR-selected loci from S cells were evaluated with composition-matched G1 and G2 cells. We then estimated the detection efficiency and corrected it for cell-to-cell batch differences. We repeated the procedure 12 times per *TSR*-bin, reporting the larger of the standard error across bootstrap repeats and the analytical GMM error. Bins containing fewer than 10,000 locus-cell pairs for E14-2025 or 500 pairs for E14-2021 were discarded as unstable.

We fitted the resulting replicated fraction as a function of *TSR* with a sigmoid *p* = *K*/ (1 + *exp*(−*B*(*t* − *M*))). The T-width was defined as the *TSR* interval over which the sigmoid transitioned from 25% to 75%. As a negative control, we randomized the S-phase cell ordering and recalculated the T-width.

### Measuring replication timing by sub-nuclear position

We measured the replicated fraction of S-phase loci occupying specific sub-nuclear positions. Given a positional criterion *L*, e.g. “loci assigned to the speckle locale”, we pooled all locus instances satisfying *L* across cells. Detection efficiency was estimated from the corresponding locus instances satisfying *L* in G1 and G2, averaged to obtain the S phase estimate, and then used to infer the replicated fraction of locus instances in S-phase cells satisfying *L*.

Subnuclear positional requirements were defined in two ways. First, within each cell loci were ranked by either local SF3A66 or Fibrillarin fluorescence intensity, or by distance to the nuclear envelope, and divided into 20 equally populated quantiles. The replicated fraction was then estimated separately for each quantile. Second, locus instances in each cell were assigned to one of eight nuclear locales. S-phase cells were ranked by their total inferred replicated fractions and divided into ten equally populated windows. Replicated fractions were then determined for all allele instances in each nuclear locale in each S-phase window. For the three sparsely populated mixed locales, four windows were used instead because of the smaller sample size.

### RepTile’s machine-learning module

The machine-learning module predicts the replication state of individual allele-resolved loci. Models were trained on loci in G1-and G2-phase cells, whose replication states were assigned as replicated and unreplicated, respectively, and then applied to loci in S-phase cells.

#### Feature extraction

For each locus allele instance and cell, we collected spot count and intensity, chromosome identity, genomic position, nuclear locale and z-coordinate. Continuous features, except spot count and genomic position, were ranked and divided into 100 quantiles; nuclear locales were one-hot encoded; and all features were smoothed using a 600kb sliding-window average.

#### Training

Separate models were trained for each chromosome. To avoid boundary effects, we excluded cells close to the G1/S and S/G2 transitions, and stratified the remaining cells into 80% training and 20% test sets, ensuring that no cell contributed to both sets. We under-sampled the majority class ensuring the same number of G1 and G2 instances, reserved 15% of balanced training data for validation, and performed standardized scaling on the training data. We trained XGBClassifier ^30^ with tree_method = ‘hist’, max_depth = 12, learning_rate = 0.05, n_estimators = 1,000, subsample = 0.8, colsample_bynode = 0.8, reg_lambda = 2.0 and reg_alpha = 1.0, stopping after 10 rounds without improved validation area under the receiver-operating-characteristic curve (AUC). The model output was the probability that a locus instance belonged to G2 and therefore was replicated, i.e. the replication probability.

#### Binarizing probabilities into states

A locus was called replicated at p ≥ X, non-replicated at p ≤ 1-X and otherwise left unassigned. Under this rule, we measured accuracy across thresholds on the G1/G2 test set. We used X = 0.9 for most analyses and X = 0.8 when greater coverage was required.

#### Performance evaluation and prediction

Model performance was evaluated on the held-out G1/G2 test set using the AUC, computed from the predicted replication probabilities. We also computed the AUC within strata of RT, z-coordinate and nuclear locale to verify that the model did not merely recapitulate a genomic or spatial detection bias. The coverage was measured as the fraction of predicted allele instances in S-phase. *Average replicated percentages.* Ignoring unassigned loci, we calculated replicated fractions as the number of replicated loci divided by all assigned loci. Averaging over cells gives locus-specific RT, while averaging over loci gives per-cell S-phase progression. For genomic correlation analysis at 25kb resolution, signals were smoothed using a 300kb sliding window.

### Single-cell Repli-seq analysis

Single-cell Repli-seq data were analyzed following the original protocol ^39^. The data provide a cell-by-locus replication-state matrix at 50kb resolution, which we mapped to the corresponding imaged bins. We calculated the S-phase progression as the replicated fraction per cell, replication timing as the replicated fraction per locus, and replication-timing heterogeneity using the T-width metric described above.

### Measuring position sensitivity of loci

To quantify how strongly a locus’s replication timing depends on its sub-nuclear position, we devised an ANOVA-style variance decomposition approach. We coarse-grained RepTile ML replication probabilities from 25kb to 200kb resolution and binarized each allele instance as replicated (P > 0.9), non-replicated (P < 0.1) or ambiguous (NA). Ambiguous instances were excluded. Each 200kb allele instance was assigned a nuclear locale by majority vote over its eight constituent 25kb sub-loci. We only analyzed autosomes, and restricted to the five primary locales (speckle, nucleoli, envelope, non-landmark active and repressive), excluding the sparsely populated mixed locales.

For each locus, we modeled its replication state *y_i_* across S-phase cell allele instances *i* as a function of the S-phase pseudo-time *T_i_* and nuclear locale *L_i_*:

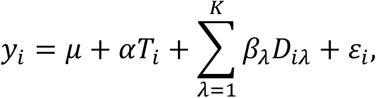

where *D_iλ_* is an indicator variable equal to 1 if instance *i* occupies locale *λ*, while *μ*, *α*, *β_λ_* are parameters to fit, and *ε* is a random variable with zero mean. Including pseudo-time accounts for differences in S-phase progression and prevents it from confounding the locale effect. We also fitted a nested reduced model without locale terms:

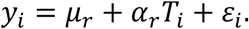

We fitted both models by ordinary least squares, yielding the best predictions *ŷ_1_* (with locales) and *ŷ_0_* (without). The difference between these predictions was interpreted as the “variation explained by locales” and quantified as ||*ŷ_1_* − *ŷ_0_*||^2^. Position-sensitivity was defined as the square root ratio between this quantity and the variance left unexplained after only regressing out S-phase pseudo-time, ||*y* − *ŷ_0_*||^2^:

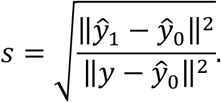

It ranged from 0, when locale carries no information, to 1, when it explains all remaining variation.

We assessed significance independently for each locus by randomly shuffling locale labels across S-phase alleles (B=10,000 times) while holding replication state and pseudo-time fixed and re-calculating position-sensitivity. P-values were calculated as

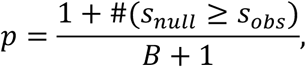

and were corrected across 200kb loci by the Benjamini-Hochberg procedure. We defined position-insensitive loci as those below the 95^th^ percentile of position-sensitivity values among loci with permutation FDR > 0.05 (s = 0.19). The remainder were position-sensitive, classified as strongly sensitive at s > 0.34 and weakly position-sensitive otherwise. The 0.34 threshold was chosen as the median plus one median absolute deviation (MAD) of the genome-wide position-sensitivity distribution.

For this ANOVA analysis, loci represented in fewer than three locales with at least ten (≥10) observations were excluded (52 of 11,872 autosomal loci). For retained loci, individual locales represented by fewer than ten observations were excluded from the locus-specific model.

### Replication propensity by nuclear locale

For each locus, we calculated the fraction of replicated allele instances in each locale *λ* across S-phase cells, *RF_i_ (λ)*. We then defined the maximum locale-dependent difference in replicated fraction as

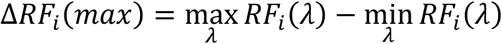

We also calculated the locale-specific replication shift of a locus *i*, defined as the difference between the replicated fraction in a given locale and that averaged across all other locales:

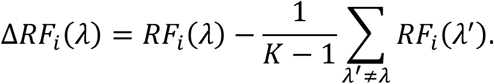

where *K*=5 is the number of primary nuclear locales. Only locales represented with ≥10 allele instances for a given locus were evaluated.

### Clustering position-sensitive loci by locale-associated replication shifts

For each position-sensitive locus, we assembled its five *ΔRF_i_ (λ)* values into a locus-by-locale matrix. Locus-locale pairs represented by fewer than 10 observations were assigned a value of 0. For each locus, we centered and scaled the *ΔRF_i_ (λ)* and applied a soft-max function with temperature *T* = 0.5:

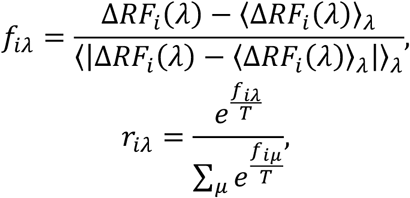

We clustered loci using the *r_iλ_* vectors with k-means (*k* = 5, 20 initializations) ^68^. The number of clusters was identified using the Kneedle algorithm ^58^. For *k* spanning from 2 to 10, both the *k* and the corresponding per-*k* inertia values, i.e. the total within-cluster sum of squared Euclidean distances, were min-max normalized, and the *k* whose point was farthest from the line connecting the normalized endpoints was selected, yielding *k* = 5.

Clusters were labeled by their ‘replication-favorable locale’ (RFL), defined as the locale showing the greatest positive replication shift: C1, speckle; C2, non-landmark active; C3, non-landmark repressive; C4, nucleoli; and C5, envelope. Within each heatmap cluster, loci were ordered by their projection onto the centroid. Position-insensitive loci were not clustered and were ordered genomically.

### Correspondence between canonical and replication favorable locale

For position-sensitive loci, we calculated, within each canonical-locale group, the percentage of loci assigned to each RFL cluster. To assess significance, we randomly shuffled each locus’s canonical locale and cluster assignment 10,000 times, preserving their marginal frequencies. Across shuffled instances and for each canonical-RFL pair, we calculated both the average matching percentage (for the observed/expected ratio) and the one-sided p-value, defined as the fraction of times a random matched percentage reached or exceeded the observed one. We corrected these p-values with the Benjamini-Hochberg procedure across the 25 canonical-RFL pairs.

We additionally calculated the overall percentage of loci for which the canonical locale matched the RFL and assessed its significance using the same permutation framework. Analyses were performed separately for strongly and weakly position-sensitive loci.

### Single-cell discordant-allele analysis

For each 200kb position-sensitive locus, we identified S-phase cells whose two alleles had opposite replication states in the same cell, with one classified as replicated (P > 0.8) and one classified as non-replicated (P < 0.2). These relaxed probability thresholds were used to increase coverage for this more stringent analysis. We retained cells in which one allele occupied the locus-specific replication-favorable locale (RFL) and the other occupied a different locale, and then recorded whether the replicated allele occupied the RFL.

### Correlation between position-sensitivity and ChIP-seq tracks

We coarse-grained chromatin immunoprecipitation sequencing (ChIP-seq) tracks by averaging their signals within each 200kb bin and computed, against the 200kb position-sensitivity signal, both the Spearman correlation and the partial Spearman correlation adjusting for canonical locales. The latter was calculated by ranking both the position-sensitivity and the ChIP-seq tracks genome-wide, subtracting from each locus the mean rank of loci sharing its canonical locale, and correlating the resulting locale-centered ranks.

### Selection of initiation zones

Because high-resolution Repli-seq is unavailable for E14, we curated conserved mESC initiation zones (IZs) by intersecting five datasets. We first identified the genomic position of candidate IZs by calling peaks in log_2_(early/late) Repli-seq data in E14 cells ^38^. Next, we intersected them with high resolution Repli-seq IZs and log_2_(early/late) peaks from hybrid (castaneus x musculus) F121-9 mESC, as well as the SNP-resolved *mus musculus* genome from the same datasets ^40^. This intersection (N=895) contains 42% of IZs across all datasets. We assigned a replication state by thresholding the average replication probability of the 25kb loci composing them, and a nuclear locale by majority vote, and then applied the position sensitivity analysis as described previously. We also generated a list of 200kb ‘non-IZ’ regions, defined as the set subtraction of all 200kb loci and the peaks and IZs from the five datasets.

### Constitutive and developmentally switching replication timing domains

Following the original protocol ^3^, we coarse-grained and quantile-normalized RT profiles from 31 mouse cell types to 200 kb. For each locus, we calculated RT_min_ and RT_max_ across cell types and classified it as constitutive early (RT_min_ > 0.3), constitutive late (RT_max_ < −0.2), developmentally switching (RT_min_ < −0.2 and RT_max_ > 0.3) or NA. The same classification was applied to the 895 IZs.

## Data availability

All datasets used in this study were downloaded from public repositories:

E14-2025 DNAseqFISH+ ^16^: https://zenodo.org/records/7693825,

E14-2021 DNAseqFISH+ ^19^: https://zenodo.org/records/3735329, mESC single-cell Repli-seq ^39^: https://www.ncbi.nlm.nih.gov/geo/query/acc.cgi?acc=GSE102077,

ES-E14TG2a with CTCF-AID two-stage Repli-seq ^38^: https://data.4dnucleome.org/experiment-set-replicates/4DNESOUOK9IO/ and https://data.4dnucleome.org/experiment-set-replicates/4DNES9G5R9GI/,

F121-9-CASTx129 LaminB1 TSA-seq: https://data.4dnucleome.org/experiment-set-replicates/4DNES1MDG5JB/,

F121-9-CASTx129 MKI67IP TSA-seq: https://data.4dnucleome.org/experiment-set-replicates/4DNES29R7A5U/,

F121-9-CASTx129 SON TSA-seq: https://data.4dnucleome.org/experiment-set-replicates/4DNESDN4HVNK/,

F121-9-CASTx129 and parsed 129 two-stage Repli-seq ^59^: https://www.ncbi.nlm.nih.gov/geo/query/acc.cgi?acc=GSE114747, F121-9-CASTx129 initiation zones ^40^: https://figshare.com/articles/dataset/Human_and_mouse_IZ_TTR_termination_sites_Twidth/25517497/2,

mESC in situ Hi-C ^34^: https://data.4dnucleome.org/experiment-set-replicates/4DNESDXUWBD9/,

E14 in situ Hi-C PC1 ^37^: https://data.4dnucleome.org/experiment-set-replicates/4DNESU4BQU4G/,

Replication Timing across 31 mouse cell types ^3,48^: https://figshare.com/articles/dataset/List_of_constitutive_and_developmentally-regulated_replication_domains_mouse_mm10_/24168963/1,

ChIP-seq datasets. From ENCODE (https://www.encodeproject.org/): H3K4me1 (ENCSR000CGN), H3K4me3 (ENCSR000CGO), H3K9ac (ENCSR000CGP), H3K27ac (ENCSR000CGQ), H3K36me3 (ENCSR000CGR), H3K27me3 (ENCSR059MBO), H3K9me3 (ENCSR000ADM). From Gene Expression Omnibus: H4K16ac ^69^ (GSE43103), Rif1 ^70^ (GSE98253), BRD4 and MED1 ^71^ (GSE112808).

## Software and code availability

Our analyses were performed in Python, using standard libraries for data analysis including *numpy*, *pandas*, *scipy*, *matplotlib*, *scikit-learn*, *statsmodels* ^72–77^. α-shapes and convex hulls were computed with the *alphashape* and *trimesh* Python libraries (https://github.com/bellockk/alphashape/tree/v1.3.1, https://github.com/mikedh/trimesh). Hi-C data were extracted and processed with the Python *cooler* package ^78^. 3D renderings of single cells were produced with *ChimeraX* ^79^. The genomic visualization in Extended Figure 7a was produced with IGV ^80^. RepTile’s code is available on GitHub:

https://github.com/alberlab/RepTile.

## Acknowledgments

This work was supported by the National Institutes of Health (NIH); grants UM1HG011593 (F.A. and D.G.) and R01GM162977 (F.A.). We thank Peiyao A. Zhao for extracting the 129 initiation zones. We thank Lorenzo Boninsegna, Ye Wang, and the rest of Frank Alber lab for insightful scientific conversations about our work.

## Author contributions

Conceptualization, F.M., D.G., and F.A.; F.M. performed all calculations, designed and performed data analysis. F.M., D.G. and F.A. interpreted results and data analysis. F.M. wrote software and documentation; writing – original draft, F.M.; writing -review and editing F.M., D.G., and F.A.; Supervision D.G. and F.A.; funding acquisition, D.G. and F.A. All authors approved the final manuscript.

