## Supplementary Figure 1 for "Decoding single-cell replication states from imaging data reveals locus-specific subnuclear position dependencies of replication initiation"

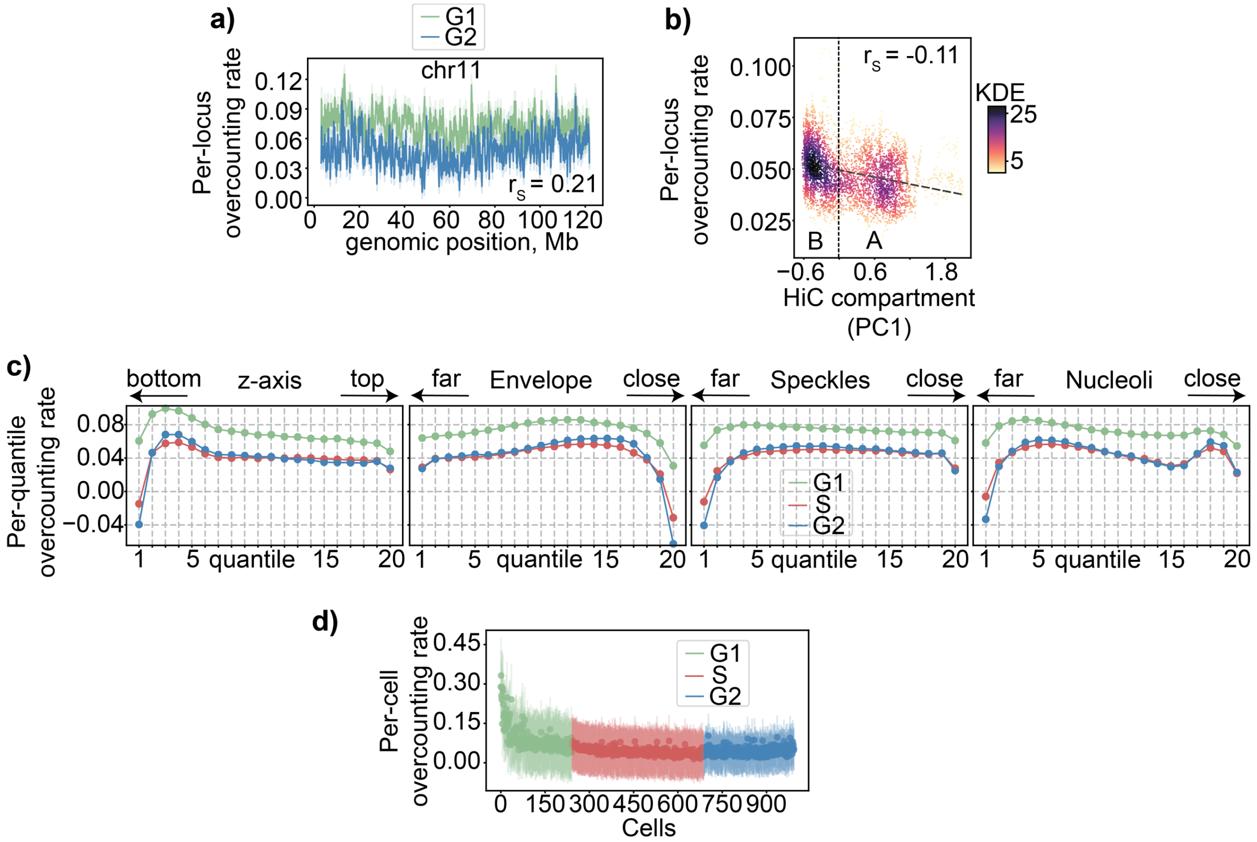


**Over-counting rate estimated by RepTile's statistical module in the E14 mouse embryonic stem cell DNAseqFISH+ 2025 dataset [1].**

**a)** Average per-locus overcounting rate for chromosome 11, inferred for G1 (green) and G2 (blue), with their Spearman correlation. **b)** Average per-locus overcounting rate plotted against Hi-C compartment PC1 [2], colored by local Gaussian kernel density estimation, with their Spearman correlation shown. The dotted vertical line indicates the separation between B (left) and A (right) compartment. The dashed line indicates the linear fit. **c)** Average per-quantile overcounting rate against 20 quantiles of z-axis, envelope distance, speckle and nucleoli association (**Methods**), separately for G1, S, G2. **d)** Average per-cell overcounting rate against cell index, with cells sorted by cell cycle progression.

1. Takei, Y. *et al.* Spatial multi-omics reveals cell-type-specific nuclear compartments. *Nature* **641**, 1037–1047 (2025).
2. Yan, J. *et al.* Histone H3 lysine 4 monomethylation modulates long-range chromatin interactions at enhancers. *Cell Res* **28**, 204–220 (2018).
